# Remembrance of things practiced: Fast and slow learning in cortical and subcortical pathways

**DOI:** 10.1101/797548

**Authors:** James M. Murray, G. Sean Escola

**Affiliations:** Zuckerman Mind, Brain, and Behavior Institute, Columbia University

## Abstract

The learning of motor skills unfolds over multiple timescales, with rapid initial gains in performance followed by a longer period in which the behavior becomes more refined, habitual, and automatized. While recent lesion and inactivation experiments have provided hints about how various brain areas might contribute to such learning, their precise roles and the neural mechanisms underlying them are not well understood. In this work, we propose neural- and circuit-level mechanisms by which motor cortex, thalamus, and striatum support such learning. In this model, the combination of fast cortical learning and slow subcortical learning gives rise to a covert learning process through which control of behavior is gradually transferred from cortical to subcortical circuits, while protecting learned behaviors that are practiced repeatedly against overwriting by future learning. Together, these results point to a new computational role for thalamus in motor learning, and, more broadly, provide a framework for understanding the neural basis of habit formation and the automatization of behavior through practice.

## Introduction

The acquisition and retention of motor skills, for example shooting a basketball or reaching for a piece of food, are crucial for humans and other animals. Such skills are acquired through repeated practice, and the effects of such practice are numerous and complex [1, 2, 3]. Most obviously, practice leads to improved performance, e.g. as measured by movement accuracy [4]. In addition, practice leads to the formation of habits, which can be defined operationally as decreased sensitivity of a behavior to goals and rewards [5, 6]. Finally, practice leads to behavioral automatization, which can be measured by improved reaction times or decreased cognitive effort [2]. In many cases, these multiple effects of practice take place on different timescales, with large gains in performance obtained early in learning, followed by a much longer period in which the learned behavior becomes more refined, habitual, and automatic [7, 8]. This suggests the possibility that motor learning can be divided into a fast, overt learning process leading to improved performance, combined with a slower, covert learning process with subtler effects related to habit formation and behavioral automatization. It is currently not well understood, however, whether these multiple learning processes that occur during practice are truly distinct or merely different facets of the same underlying process, whether they operate sequentially or in parallel, and how they might be implemented in the brain.

Neurobiologically, the descending pathway through the sensorimotor region of the striatum, the major input nucleus of the basal ganglia, plays an important role in both the learning and execution of learned motor behaviors [9, 10, 11], as well as in habit formation for highly practiced behaviors [12, 13]. Such learning is facilitated by dopamine-dependent plasticity at input synapses to striatum, where this plasticity plays a role in reinforcing behaviors that have previously led to reward [14]. The sensorimotor striatum is able to influence behavior through two pathways: one descending pathway to motor-related brainstem nuclei [15] and another pathway to motor cortex via the basal ganglia output nuclei and motor thalamus [16]. Neural activity in sensorimotor striatum itself is driven by input from both cortical and subcortical sources. The dominant cortical input comes from sensorimotor cortex [17, 18], and a great deal of research has implicated this pathway in motor learning [19, 20]. The dominant subcortical input, meanwhile, comes from thalamus, including the rostral intralaminar nuclei and parafascicular nucleus [21, 22]. Although this subcortical input to striatum has received less attention than the corticostriatal pathway, it has also been shown to represent behaviorally relevant variables [23, 24, 25] and to be important for the production of learned behaviors [26]. In general, however, the role of this thalamic pathway in motor learning and how it might differ from the corticostriatal pathway is not well understood.

In order to obtain some understanding about the roles that cortex, thalamus, and striatum play in motor learning, experimental manipulations of these structures and of the projections between them during different stages of learning provide useful constraints. Recent lesion and inactivation experiments in rats have shown that sensorimotor striatum and its thalamic inputs are necessary for both learning and execution of a skilled lever-press sequence [27]. In the same task, however, input from motor cortex to striatum is necessary for learning the task, but not for performing the task once it has already been learned [28, 27]. Furthermore, the representation of task-related kinematic variables in striatum during the task is similar in rats with and without cortical lesions [29]. Evidence for the gradual disengagement of motor cortex during the learning of skilled movements has also been found in mice [30] and humans [31, 32]. Together, the above results provide hints about the roles played by cortex, thalamus, and striatum in the acquisition and execution of skilled movements, suggesting that responsibility for driving learned behaviors may be gradually offloaded from cortical to subcortical circuits through practice. It is not immediately obvious, however, which neural- or circuit-level mechanisms might explain these results, nor how they might relate to the multiple effects of practice discussed above.

In this work, we propose a mechanistic theory of learning at cortico- and thalamostriatal synapses that provides a unified description of the lesion and inactivation results described above and, more broadly, points to computationally distinct roles for the two inputs to striatum in the acquisition and retention of motor skills. As a starting point for addressing learning, retention, and recall accuracy, we begin by modeling a single neuron within striatum. We mathematically derive the neuron’s forgetting curve, which quantifies the probability that the neuron produces the correct output in response to a particular pattern of inputs as a function of how far in the past learning occurred. If the neuron’s input synapses are all modified by a supervised (i.e. error-minimizing) plasticity rule, then the neuron learns to produce the correct output in response to a particular input, but there is no added benefit to repeated practice once the output has been learned. If, on the other hand, some of the neuron’s input synapses are modified by a slower, associative learning rule, then patterns that are practiced many times become far more resistant to being overwritten by future learning. We interpret the first input, with fast supervised learning, as motor cortex, and the second input, with slow associative learning, as thalamus. We further show that, as a behavior is practiced, there is a gradual and covert transfer of control, with the activity of the downstream population increasingly driven by the second input pathway. In addition to providing a unified description of the lesion and inactivation experiments described above, this model generates predictions for the expected effects of perturbations to the cortical and thalamic pathways. Further, it proposes a new computational role for thalamic inputs to striatum and, more broadly, provides a framework for understanding habit formation and the automatization of behavior as the control of behavior is transferred from cortical to subcortical circuits.

## Results

In the following sections, our aim will be to develop a theory that describes the effects of learning and practiced repetition described above. As a first step, we will mathematically investigate learning and forgetting within a simple model of supervised learning in a single striatal neuron receiving cortical inputs. This will provide a minimal quantitative theory for addressing questions about retention and accuracy in learned behaviors, but will not yet account for effects related to practiced repetition. In order to account for such effects as well as for the lesion and inactivation results described above, we will next add a second set of inputs from thalamus. By endowing these inputs with associative learning, we will show that this second input pathway leads to enhanced retention and greater accuracy for patterns that are practiced many times during training. These two types of learning together give rise to a covert learning process in which control of the striatal population is gradually transferred from the cortical to the thalamic inputs through repeated practice. Finally, we describe how this two-pathway model leads to a number of experimental predictions about the expected effects of perturbations and inactivations of the cortical and thalamic inputs.

### A single-neuron model describes learning and forgetting

In order to develop a quantitative theory of learning at the most basic level, we began by studying a classical model of a single neuron: the perceptron [33], which in our model corresponds to a single striatal neuron receiving cortical input (Figure 1a,b). In this model, *N*_*x*_ input signals *x*_*i*_ are multiplied by synaptic weights *w*_*i*_ and then summed. If the summed value is greater than 0, the neuron’s output is +1, otherwise the output is −1. This is described by the equation *z*^*µ*^ = sgn(**w**^*µ*^ · **x**^*µ*^), where *µ* denotes one of *P* possible input patterns (Figure 1b). The goal is then for the synaptic weights **w** to be adjusted so that, for each pattern *µ*, the output matches a target value 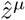. Thus, the task is binary classification, in which random input patterns (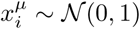, where 𝒩 denotes the standard normal distribution) are mapped onto random output values 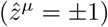. In our neurobiological interpretation, the perceptron corresponds to a single neuron in striatum, the major input structure of the basal ganglia. This neuron receives inputs from cortex with information about cues, context, and sensory feedback, and it learns to adjust the strengths of its input weights so that the neuron is active in response to some inputs and inactive in response to others.

**Figure 1:**
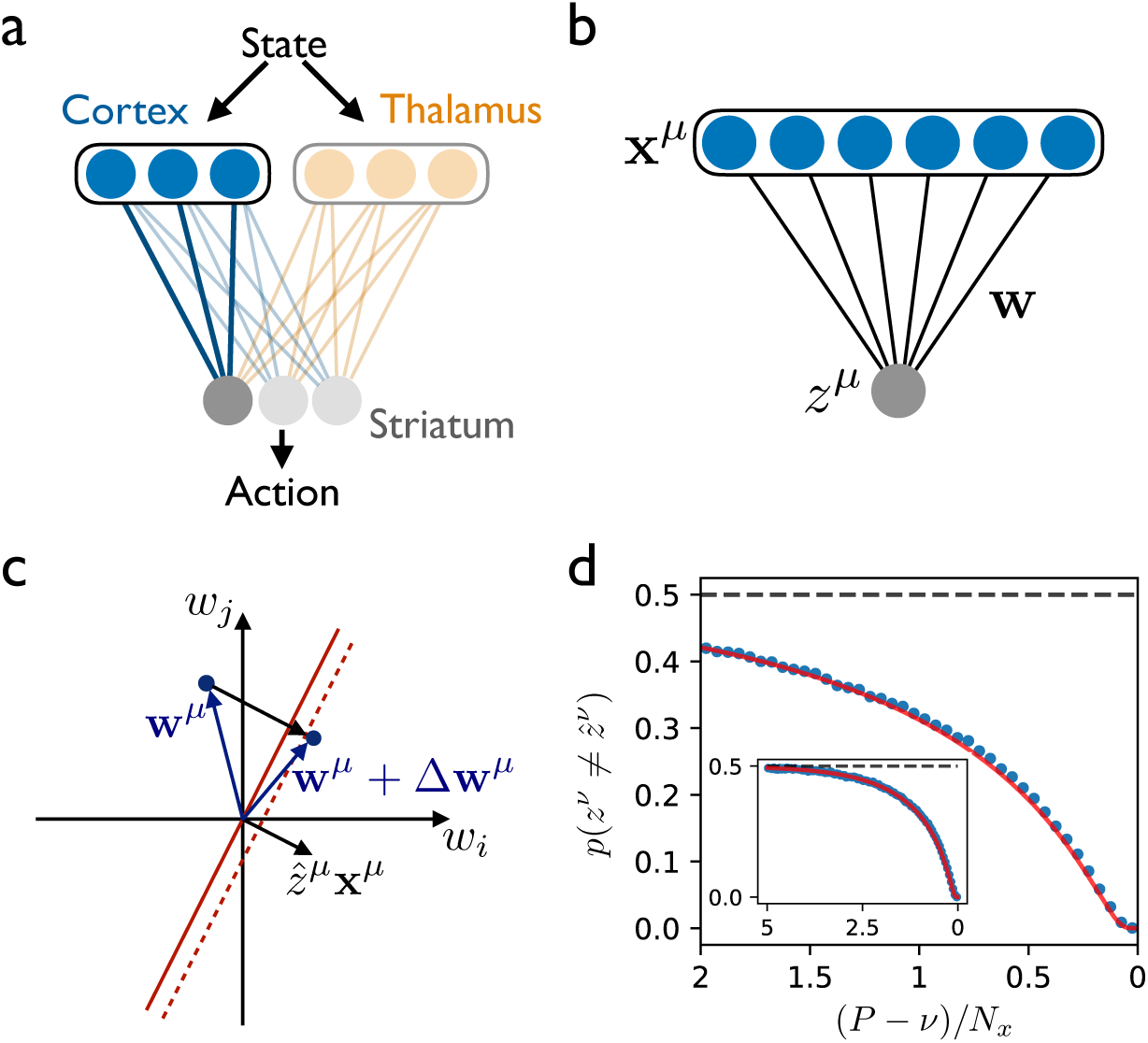
The forgetting curve for a single neuron with supervised learning. **(a)** A sensorimotor circuit model in which cortex and thalamus contain state information (e.g. cues and sensory feedback) and provide input to striatum, which learns to drive actions that lead to reward. **(b)** A model of a single striatal neuron receiving cortical inputs, where weights **w** are trained such that random input patterns **x**^*µ*^ produce the correct classifications 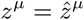. **(c)** The input vector **x**^*µ*^ defines a hyperplane in the space of weights **w**, and the update rule modifies the weight vector to give the correct classification with margin *κ* = 1. **(d)** The probability of incorrect classification when testing pattern *ν* after learning *P* = 2*N*_*x*_ patterns sequentially. The most recently learned patterns are on the right, with earlier patterns on the left. The solid curve is the theoretical result; points are simulated results. Inset shows the same result for a perceptron trained with *P* = 5*N*_*x*_ patterns.

This learning problem can be illustrated geometrically in the *N*_*x*_-dimensional space of synaptic weights, where the weight vector **w**^*µ*^ just before learning pattern *µ* is represented as a vector in this space (Figure 1c). In order for the classification to be correct, the weight vector after training should have positive overlap with the vector 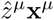, which defines a classification boundary (red line in Figure 1c), with all weight vectors on one side of this boundary leading to correct classification. This can be effected by updating the weights to **w**^*µ*^ + Δ**w**^*µ*^, where

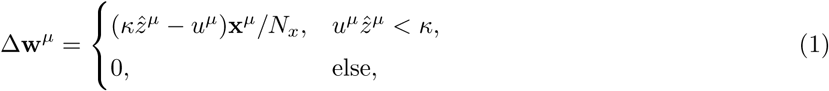

where *u*^*µ*^ = **w**^*µ*^ · **x**^*µ*^ is the summed input to the neuron [34]. The first line of this equation says that, if the classification is initially incorrect or correct but within a margin *κ* of the classification boundary (dashed line in Figure 1b), then an update is made such that the new weight vector **w**^*µ*^ + Δ**w**^*µ*^ lies on the correct side of the classification boundary (Figure 1c) with margin *κ*. The second line, on the other hand, says that, if the classification is initially correct with a margin of at least *κ*, then no update needs to be made. Because *κ* can be absorbed into an overall rescaling of **w**, we henceforth set *κ* = 1.

Clearly a potential problem exists whenever more than one pattern is being learned, however. After applying (1) to ensure correct classification of pattern *µ*, followed by applying the same update rule for pattern *µ* + 1, there is no guarantee that the weight vector will still be on the correct side of the classification boundary for pattern *µ*. That is, the learning of new information in this model has the potential to interfere with and overwrite previously learned information. In the standard paradigm for learning in the perceptron, this problem is solved by cycling many times through all of the *P* patterns to be learned. In one of the most celebrated results in computational neuroscience, it has been shown that, in the large-*N* limit, a weight vector producing correct classification of all patterns can be obtained whenever *P <* 2*N*_*x*_ [35, 36].

In order to address effects related to retention and recall, we instead study the case in which the patterns are learned in sequence, so that no further training with pattern *µ* occurs after training with pattern *µ*+1 has begun. Concretely, we first train the perceptron by applying the update rule (1) for each of the *P* patterns in the sequence. We then test the perceptron by testing the classification of each pattern using the final weight vector **w**^*P*^. Intuitively, we expect that the most recently learned classifications will remain correct with high probability during testing since the weight vector will have been updated only a small number of times after learning these patterns. On the other hand, for classifications learned much earlier and hence overwritten by a large number of subsequent patterns, the probability of incorrect classification should approach chance level.

In order to describe this process of training, overwriting, and testing quantitatively, we calculated the probability of incorrect classification (i.e. the error rate) for each pattern during testing after all patterns have been trained. The full calculation is presented in Appendix 1. To summarize, it begins by assuming the weight vector is **w**^*ν*^ just before learning some generic pattern *ν*, then applies the update rule (1) using randomly drawn **x**^*ν*^ and 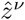 to obtain Δ**w**^*ν*^. For the following *P* − *ν* steps (*µ* = *ν* + 1, …, *P*), the updates Δ**w**^*µ*^ will be in directions that are random with respect to **w**^*ν*^. Hence, the evolution of the weight vector from **w**^*ν*^ to **w**^*P*^ can be described as a drift-diffusion process characterized by the probability distribution *p*(**w**^*P*^ |**w**^*ν*^ + Δ**w**^*ν*^). Once the probability distribution for **w**^*P*^ is known, the error rate during testing is calculated by averaging *z*^*ν*^ = sgn(**w**^*P*^ · **x**^*ν*^) over **w**^*P*^, as well as over the initial weight vector **w**^*ν*^ and the random input pattern **x**^*ν*^.

The result of this calculation is that the error rate during testing of pattern *ν* is given by

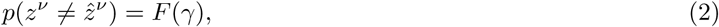

where the form of the function *F* is given in Appendix 1, and 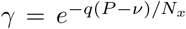, where *q* ≃ 0.798 is the probability that a nonzero update is made according to (1) during any particular step of training. This result defines the single-neuron forgetting curve, which is shown in Figure 1d. (While the pattern index *ν* is the independent variable in the forgetting curve, we plot the curve as a function of (*P* − *ν*)*/N*_*x*_ so that earlier times appear on the left and later times on the right, and so that the curve is independent of *P*, which may be arbitrarily large.) As expected, the forgetting curve in Figure 1d shows that recently learned classifications are recalled with very low error rates, while those learned far in the past are recalled with error rates approaching chance level. We can further observe that, because the number of synapses *N*_*x*_ appears only through the parameter *γ*, the number of patterns that can be learned with error rate at or below any particular threshold is proportional to *N*_*x*_. This means that the memory is extensive in the number of synapses, as in the classical result described above for the perceptron with cycled training. Finally, because the perceptron implements learning in a single step and may therefore be questionable as a neurobiological model of learning, we show using simulations that a similar forgetting curve is obtained using a gradient-descent learning rule, in which small updates are accumulated over many repetitions to minimize the readout error (Supplemental Figure 1).

The single-neuron model that we have studied in this section provides a zero-parameter baseline theory for addressing questions related to learning, retention, and recall accuracy. The ability of this model to learn new information by overwriting recently learned information is in contrast to the case of the classical perceptron, in which patterns are cycled through repeatedly during training, leading to a classification error rate of zero for up to *P* = 2*N*_*x*_ patterns. However, no further classifications could be learned in this case without drastically impairing the classification performance of the already-learned patterns. This phenomenon, known as catastrophic forgetting [37], is obviated in the sequentially trained model studied here. As previously pointed out in theoretical studies of related models [38, 39], although sequentially trained neurons and neural networks have a smaller overall memory capacity than models in which training is repeated, they have the advantage that learned patterns decay smoothly over time as new patterns are learned, even as *P/N*_*x*_ → ∞.

### A two-pathway model describes the effects of practice

In the preceding section, we derived the forgetting curve for a single-neuron model of a striatal neuron receiving cortical inputs with supervised learning in order to quantitatively address learning and forgetting. Because learning of any given pattern occurs in a single step and stops once the error is zero, however, this simple model is unable to describe the effects of practice. In order to study the effects of practice, we made the simplest possible addition to the model by including a second input pathway, where synaptic weights in this pathway are modified with an associative update rule, as originally proposed by Hebb [40] (Figure 2a). Specifically, these synapses are strengthened by coactivation of their inputs and outputs, so that the same inputs will in the future tend to produce the same outputs. This strengthening, which is independent of errors or rewards, continues to accumulate with each repetition if an input pattern is presented many times. Importantly, the Hebbian learning should be slow compared to the supervised learning, so that it will not strengthen an incorrect input-output association before the supervised learning has taken place. Intuitively, then, this picture suggests that repeated training of a particular classification should strengthen the association and make it more resistant to being overwritten by later learning. We shall show below that this is indeed the case. After developing the theory for this two-pathway model, we shall show in the following section that identifying the second input to striatum as thalamus provides an explanation for the asymmetric effects of motor cortical and thalamic lesions in rats described in the Introduction [28, 29, 27].

**Figure 2:**
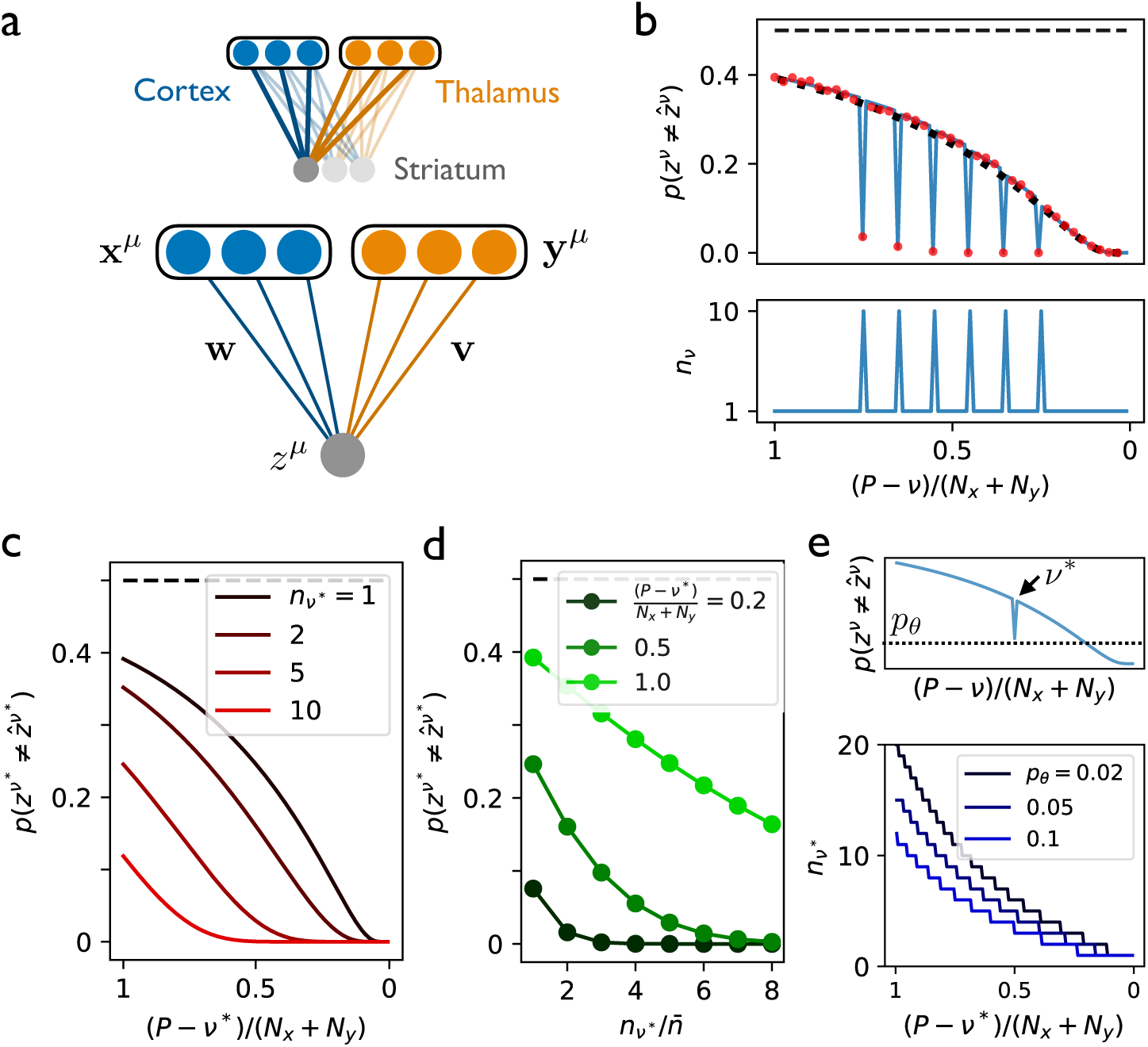
Repetition during training improves recall in the two-pathway model. **(a)** The two-pathway architecture, with cortical and thalamic inputs to striatum (*top*), where fast supervised learning of corticostriatal weights **w** is accompanied by slow Hebbian learning of thalamostriatal weights **v** (*bottom*). **(b)** The forgetting curve (*top*) in a case where six patterns are repeated multiple times during training, while all other patterns are presented only once (*bottom*). Solid curve is theoretical result (*α* = 1, *β* = 1); points are simulations with *N*_*x*_ = *N*_*y*_ = 1000; dotted curve shows the case in which no patterns are repeated multiple times. **(c)** The error rate for pattern *ν*\*, which is repeated *n*_*ν*\*_ times during training, while other patterns are presented only once. **(d)** The error rate as a function of *n*_*ν*\*_, with curves corresponding to different choices of *ν*\*. **(e)** *Top*: A single pattern *ν*\* is trained multiple times while all other patterns are trained once. *Bottom*: the number of times that pattern *ν*\* must be repeated during training in order to obtain a classification error rate below a threshold *p*_*θ*_ during testing.

Concretely, we extended the single-neuron model by adding a second set of inputs **y**^*µ*^, where the *N*_*y*_ inputs 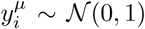 are random and independent of the first set of *N*_*x*_ inputs 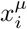. The output is then given by *z*^*µ*^ = sgn(**w** · **x**^*µ*^ + **v** · **y**^*µ*^), where the synaptic weights **v** are updated according to a Hebbian learning rule. Specifically, if **v**^*µ*^ is the weight vector just before training on pattern *µ*, then the weight vector just after training on pattern *µ* is given by **v**^*µ*^ + Δ**v**^*µ*^, where

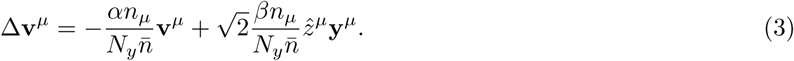

The second term in this equation is a modification to the synaptic weight that is proportional to the product of the pre- and post-synaptic activity, making this a Hebbian learning rule. Because the learning of **w** is assumed to be fast relative to this Hebbian learning, the downstream activity will have attained its target value by the time the Hebbian learning has any significant effect, allowing us to use 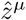 rather than *z*^*µ*^ in (3). The *n*_*µ*_ appearing in (3) is interpreted as the number of times that the behavior *µ* is repeated when it is trained, and 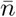 is the average of *n*_*µ*_ over all patterns. The first term in (3) causes the weights to decrease slightly in magnitude with each repetition of each pattern, so that they do not become arbitrarily large after a large number of classifications have been learned. The constants *α* and *β* control how quickly the weights decay and how strong the initial updates are, respectively. The weights **w**, meanwhile, are updated as before, except that now the total summed input appearing in (1) is given by *u*^*µ*^ = **w** · **x**^*µ*^ + **v** · **y**^*µ*^. In our neurobiological interpretation, the second pathway corresponds to thalamic inputs to the striatal neuron. As suggested by experiments [23, 24, 25], these inputs, like cortical inputs, may convey task-relevant information about cues and sensory feedback. Unlike the cortical inputs, however, the thalamic inputs connect to the readout neuron with synapses that are modified by an associative update rule.

For this two-pathway model, we derived a generalized forgetting curve using the same approach as that used for the perceptron, describing the evolution of both weight vectors **w** and **v** as drift-diffusion processes and computing statistical averages over weights and patterns. The generalized forgetting curve for the two-pathway model has the form

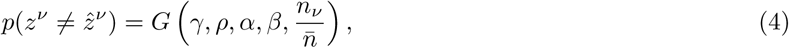

where 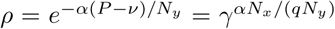, and the function *G* is derived in Appendix 2. From (4), we see that the forgetting curve now additionally depends on the Hebbian learning rate *β*, the ratio *N*_*y*_*/N*_*x*_, the Hebbian decay rate *α*, and the number of repetitions *n*_*ν*_ for each pattern.

In Supplemental Figures 2 and 3, we fixed *n*_*ν*_ to be constant for all patterns and investigated the effects of *α, β*, and *N*_*y*_*/N*_*x*_ on the forgetting curves. We found that, while adding the second pathway does not in general improve the memory performance per synapse (of which there are *N*_*x*_ + *N*_*y*_), it does improve the memory performance per *supervised* synapse (of which there are *N*_*x*_). Further, when the Hebbian decay rate *α* becomes small, the number of very old patterns that can be correctly recalled is improved, even if normalizing by the total number of input synapses (i.e. by *N*_*x*_ + *N*_*y*_, rather than by *N*_*x*_).

We next investigated the case in which some patterns are repeated more than others during training. In this case, we found that repeating particular patterns consecutively during training causes those patterns to be retained with lower error rates and for much longer than nonrepeated patterns. If trained with enough repetitions, particular patterns can be recalled essentially perfectly even after ∼ (*N*_*x*_+*N*_*y*_) additional patterns have been learned (Figure 2b). Further, as long as the number of highly practiced patterns is much less than *N*_*x*_ + *N*_*y*_, the error rate for the remaining patterns is not significantly increased (solid line versus dotted line in Figure 2b). This shows that it is possible for a neuron to become an “expert” at a small number of classifications without performing significantly worse on other classifications that have recently been learned. We next investigated the dependence of the error rate for a particular pattern *ν*\* on the interval (the “testing interval”) between training and testing, as well as on the number of repetitions *n*_*ν*\*_, for that pattern, again assuming that all other patterns are presented only once during training. We found that the effect of practice is to shift the forgetting curves in a roughly parallel manner (Figure 2c). This parallel shifting of forgetting curves via practiced repetition is a well known result from experimental psychology studies in human subjects performing memory tasks [41, 42, 43, 44, 45]. Plotting the data instead as a function of the number of repetitions *n*_*ν*\*_ for a fixed testing interval shows the error rate smoothly decreasing with practice (Figure 2d), again bearing similarity to results from memory studies in humans [46]. This shows that the error rate during testing, for a fixed recency of the practiced pattern, decreases as a function of how many times that pattern was practiced. We next asked, if just a single pattern *ν*\* is repeated multiple times while all other patterns are trained just once, how many times must pattern *ν*\* be repeated in order to obtain an error rate below a threshold *p*_*θ*_ during later testing of that pattern? The number of necessary repetitions was found to be a supralinear function of the interval between the training and testing of pattern *ν*\* (Figure 2e). This dependence could potentially be (and, to our knowledge, has not already been) measured experimentally, for example in human subjects learning paired associations (as in, e.g., Refs. [44, 45]), with certain associations presented multiple times in succession during training, and with varying intervals between training on these repeated patterns and testing.

### Input alignment and control transfer in the two-pathway model

In the previous section, we showed that, due to the reinforcement of the input-to-output mapping that comes about by slow Hebbian learning in the single-neuron model, input from the second pathway can contribute to driving the readout unit to its target state. We next investigated how the learning process in the two-pathway model could be interpreted at the population level. In this case, while each individual readout neuron performs the same binary classification task as before, the activity of the readout population is characterized by an *N*_*z*_-dimensional vector **z**. Interpreting the two input populations as cortical and thalamic inputs to striatum, we asked what are the principles that determine the relationship of this striatal population activity to its inputs, and how does this relationship evolve through learning. To address this, we defined **m** and **h** as the *N*_*z*_-dimensional inputs to the readout population from the first and second pathway, respectively (Figure 3a), where 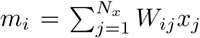 and 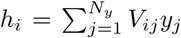, and the weight vectors **w** and **v** from the single-neuron model have been promoted to weight matrices **W** and **V**. The activity in the readout population is then described by the *N*_*z*_-dimensional vector **z** = sgn(**m** + **h**). In this case, applying the updates (1) and (3) for each synapse when training on pattern *µ* leads to updated input currents 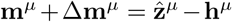 and 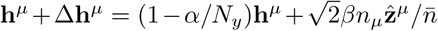 in the two pathways. Thus, following the updates, both of the input currents obtain a component of magnitude ∼ *O*(1) along the target activity 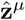, so that they become aligned with the target activity and with each other. Further, for the Hebbian update, this component continues to grow as the pattern is repeated *n*_*µ*_ times in succession (Figure 3b).

**Figure 3:**
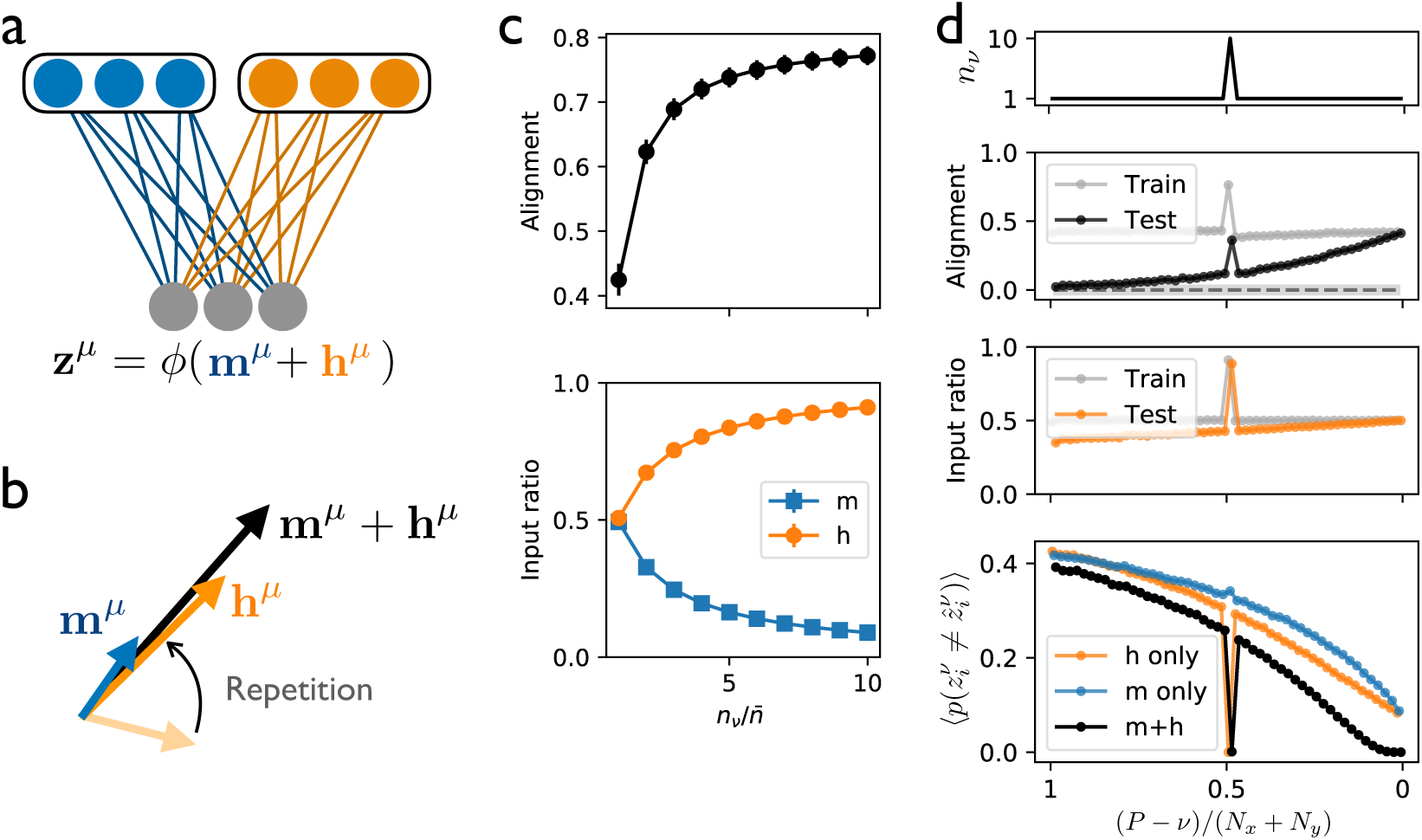
Inputs to the downstream population become aligned through repetition. **(a)** A downstream population receives input from two pathways, where the first input **m** is trained with fast supervised learning, while the second input **h** is modified with slow Hebbian learning. **(b)** Through repetition, the second input becomes increasingly aligned with the first, while its projection along the readout direction 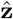 grows. **(c)** *Top*: The normalized overlap between **m** and **h** for one particular pattern that is repeated *n*_*ν*_ times. *Bottom*: The ratio of the projection of each input component along the readout direction (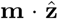 or 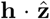) to the total input along the readout direction 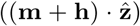. **(d)** *Top*: In a two-pathway network trained on *P* patterns sequentially, one pattern is repeated 10 times during training, while others are trained only once. *Middle panels*: The alignment for each pattern during sequential training and testing (upper panel, where shaded region is chance alignment for randomly oriented vectors), and the ratio of the input **h** along 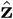 to the total input along 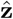 (lower panel). *Bottom*: The error rate, averaged over *N*_*z*_ readout units, for the trained network with either both inputs intact (black) or with one input removed (blue and orange) (curves offset slightly for clarity). All points in (c)-(d) are simulations with *α* = *β* = 1 and *N*_*x*_ = *N*_*y*_ = *N*_*z*_ = 1000.

As a single input pattern is repeated more times in succession, the normalized alignment between the inputs, defined as **m** · **h***/*(|**m**||**h**|) grows, meaning that the inputs from the two pathways are driving the downstream population in similar ways (Figure 3c, top), a process that we refer to as *input alignment*. In addition, as the number of repetitions grows, the input from the second pathway along 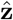 constitutes an increasingly dominant proportion of the total input along 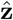 (Figure 3c, bottom), a process that we refer to as *control transfer*. Both of these processes—alignment of the two inputs and the transfer of control from one pathway to the other—are *covert*, in the sense that they occur gradually during repetition and do not have any immediately obvious behavioral correlate, since the learning in the first pathway causes the activity in the downstream population to attain its target value already by the first repetition.

In order to further illustrate the implications of input alignment and control transfer, we sequentially trained a two-pathway network with a multi-neuron readout population to produce *P* classifications, generalizing the task from previous sections to the case *N*_*z*_ *>* 1. During training, one particular pattern was repeated many times in succession, while all others were trained only once (Figure 3d, top). We found that input alignment became large and positive during training and, particularly for the repeated pattern, remained large and positive during testing after training was complete (Figure 3d, second panel). Similarly, the ratio of the input **h** along 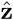 from the second pathway to the total input along 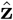 became large for the repeated pattern during training and remained large during testing, illustrating the occurrence of control transfer (Figure 3d, third panel). Finally, when testing the error rate for each pattern after training was complete, we performed “lesions” by shutting off each of the inputs during testing (Figure 3d, bottom). In the case **m** = 0, we observed that the error rate increased for all patterns *except* for the pattern that was repeated multiple times during training, which retained an error rate of zero. This shows that, via the mechanisms of input alignment and control transfer for the highly practiced behavior, the second pathway was able to compensate for the loss of the first pathway, driving the downstream population to produce the correct activity even in the absence of the first input. In the case **h** = 0, on the other hand, the repeated pattern was not recalled accurately, illustrating that the input from the second pathway is necessary to protect practiced patterns from being overwritten. Thus, interpreting the two input pathways as cortical and thalamic inputs to striatum, the model is able to account for the experimental observations in rodents that skilled behaviors and neural representations of kinematic variables are robust to the loss of cortical inputs [28, 30, 29], but that such behaviors are not robust to the loss of thalamic input [27].

In summary, when an input pathway with fast supervised learning and another with slow associative learning together drive a downstream population, the mechanisms of input alignment and control transfer enable the second input to gradually take over the role of the first in driving the downstream activity. For highly practiced patterns, this enables the second input to produce the correct downstream activity even if the first input is removed completely. As shown in Supplemental Figure 6, these mechanisms also occur in the case where reinforcement learning is used rather than supervised learning in the first pathway, but not in the case where both pathways feature reward-driven learning, illustrating that the combination of Hebbian learning together with error- or reward-based learning is necessary in order for input alignment and control transfer to take place.

### Implementation in the sensorimotor brain circuit: Automatization of behavior through practice

In the previous section, we showed that, for a single-layer neural network in which two input pathways operate with different types of learning, fast error-driven learning in the first pathway causes the downstream neurons to produce the correct output, while slow associative learning in the second pathway reinforces the association between the input to the network and the correct output. After many repetitions, this scheme enables the second input pathway to assume control for driving the downstream layer. In this section, we develop the experimental interpretation of the model in greater detail, using it to address existing experimental results and to formulate predictions for future experiments.

As a neurobiological implementation of the model, we proposed identifying the readout population in the two-pathway model as sensorimotor striatum, the input population with fast supervised learning as motor cortex, and the input population with slow associative learning as thalamus. Taken together, these identifications suggest a picture in which information about state, including sensory feedback, cues, and context, is represented in cortex and thalamus, and in which plasticity at the synapses descending from these structures to striatum allows for state information to be mapped onto actions that will tend to minimize errors and maximize rewards (Figure 4a). The mechanisms and roles of learning in the two pathways are distinct, with fast changes to corticostriatal synapses serving to learn and produce the correct behavior, while slower changes to thalamostriatal synapses serve to associate a given state with a behavioral output, with this association becoming stronger and more persistent the more that it is practiced. As described in the previous section, the transfer of control from corticostriatal to thalamostriatal inputs would suggest that motor cortex should be necessary for the initial learning of a behavior, but not for the production of the behavior after a sufficient amount of practice. This is consistent with lesion and inactivation studies in rodents [28, 30], as well as with studies of learned motor behaviors in humans suggesting that motor cortex becomes less involved in driving behaviors with practice [31, 32].

**Figure 4:**
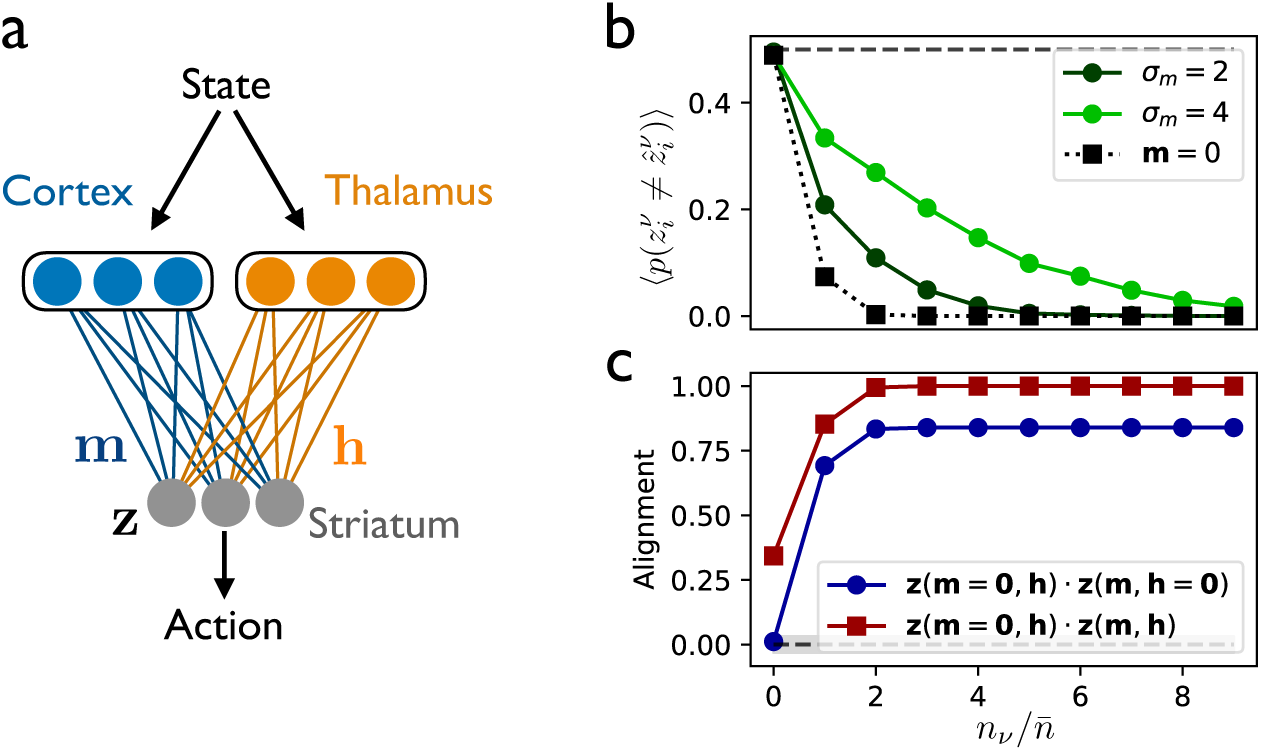
Two-pathway implementation in the sensorimotor brain circuit and predicted responses to perturbations. **(a)** In the proposed neurobiological implementation of the two-pathway model, motor cortex and thalamus provide inputs to the sensorimotor region of striatum. **(b)** The population-averaged error rate in the readout population in the presence of noise perturbations to **m** (noise amplitude *σ*_*m*_, green lines) or upon removing the input **m** (dotted line) for a single pattern repeated *n* times during training. **(c)** The normalized alignment between the population activity **z** with **m** = **0** and with **h** = **0** (blue line), and the normalized alignment between **z** with **m** = **0** and with **m** ≠ **0**. In (b) and (c), points are from simulations in a network with *α* = *β* = 1 and *N*_*x*_ = *N*_*y*_ = *N*_*z*_ = 1000.

With the neurobiological implementation proposed above, the two-pathway model also leads to a number of testable experimental predictions. First and most obviously, the theory predicts that cortico-vs. thalamostriatal synapses should be modified by different types of plasticity (fast and reward-driven vs. slow and associative, respectively). Because both synapses are glutamatergic, they have typically not been studied separately or even distinguished in past experiments [47]. Future experiments could test for differences between the types of modifications at these synapses, either *in vivo* or in slice preparations.

Second, because motor cortex plays less of a role in driving striatum as behaviors become highly practiced, lesioning, inactivating, or randomly perturbing activity in motor cortex during behavior should have a decreasingly small effect on a learned behavior the more that it has been trained (Figure 4b). Furthermore, due to the fact that covert learning in the second pathway should occur slowly relative to learning in the first pathway, the learned behavior should become robust to motor-cortex inactivation only some time after the behavior has been mastered. This effect has in fact already been observed in very recent experiments using optogenetic inactivation throughout learning of a targeted forelimb task [30].

Third, online optogenetic inactivation could be used so that only motor cortex or only thalamus drives striatum in a given trial. The theory would predict that the striatal activities observed in these two cases would be correlated due to input alignment, with the degree of correlation increasing with the amount of training (Figure 4c, blue line). Similarly, striatal population activity should be similar before and after cortical lesion or inactivation in a trained animal, with the degree of similarity increasing with the amount of training (Figure 4c, red line).

Finally, if thalamostriatal plasticity is blocked during training of a behavior, then the model predicts that the behavior will still be learned due to the fact that motor cortex is responsible for driving initial learning. Since input alignment cannot occur in the absence of thalamostriatal plasticity, however, the learned behavior will no longer be robust to cortical lesion or inactivation. This would be a challenging experiment to perform due to the fact that both thalamo- and corticostriatal synapses are glutamatergic, making selective manipulation difficult. If it could be done, however, it would provide strong evidence for our proposed neurobiological implementation of the two-pathway model.

### Simulated reaching task with the two-pathway network

With the experimental interpretation from the preceding section in mind, we asked whether the basic mechanism illustrated in Figure 2—namely, the enhanced retention of learned classifications that are practiced multiple times—could also be implemented in a neural network performing a more challenging sensorimotor task. We thus extended the two-pathway architecture to a neural network with one hidden layer and trained the network to perform a center-out reaching task (Figure 5a,b). In this task, the velocity of a cursor moving in two dimensions is controlled by a neural network, which receives as input the current location of the cursor together with a context input indicating which target to move toward. In each trial, the network was trained with supervised learning to follow a smooth, minimal-jerk trajectory to the specified target, with each trial having a duration of 10 timesteps. As in the single-neuron two-pathway model, the inputs were divided into two pathways, with supervised learning in one pathway and Hebbian learning in the other.

**Figure 5:**
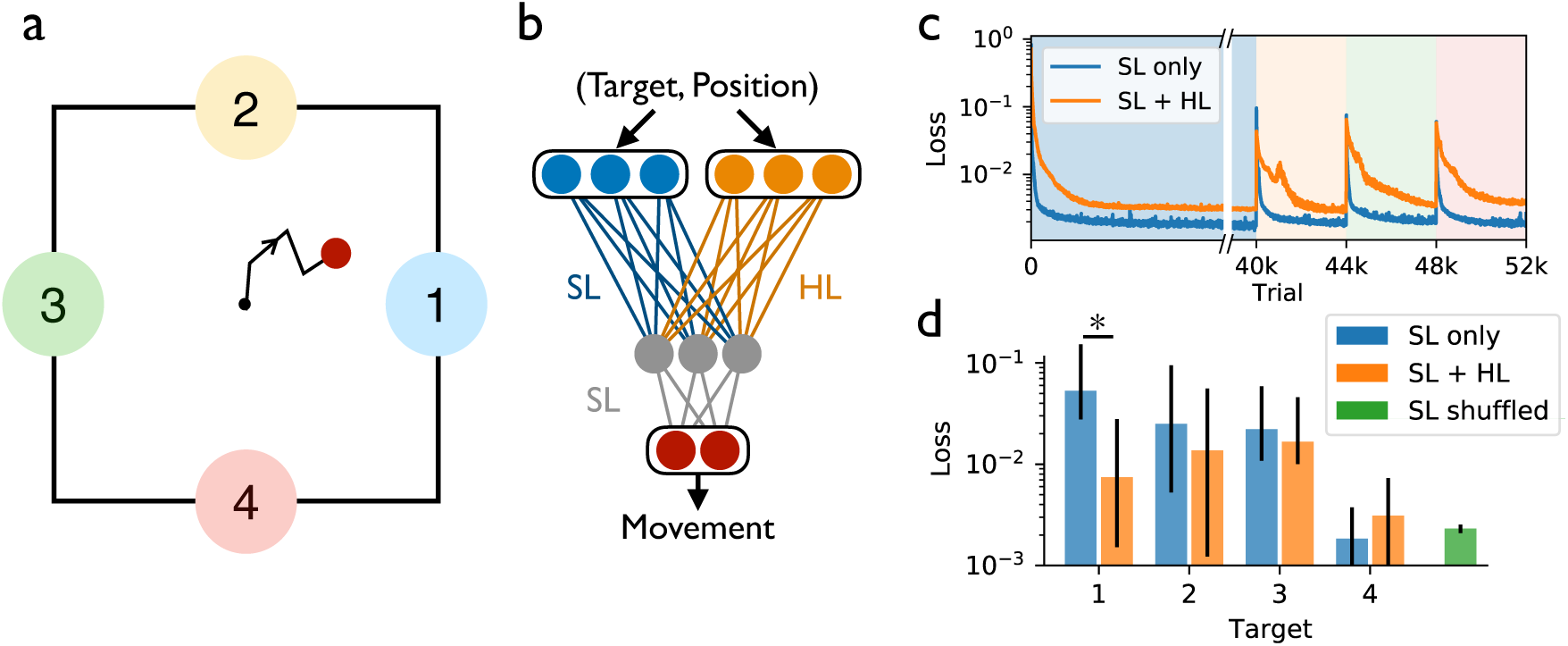
A two-pathway neural network trained to perform a reaching task exhibits enhanced retention of highly practiced behaviors. **(a)** In this task, the agent, given one of four cues, is trained to perform a center-out reach over 10 timesteps to a particular target. **(b)** The neural network trained to perform the task receives a target-specific cue and the current position as input. The network contains one pathway learned with supervised learning (SL) and another with Hebbian learning (HL). The output is a two-dimensional vector determining the movement velocity in the next timestep. **(c)** The loss (mean squared error) on each of the four targets in sequence during training, with a longer training period for the first target, and with Hebbian learning either active (SL+HL) or inactive (SL only). The loss during testing on each of the four targets, after training is complete. The green bar shows a control network in which targets were randomly interleaved during training.

The network was first trained in a block of trials to reach to target 1, then in another block to reach to target 2, and so on, with more trials in the first block than in the later blocks (Figure 5). When the network was tested after training was complete, we found that, when Hebbian learning was activated, the performance for target 1 was comparable to or even better than that for target 2, despite the fact that the training for target 2 had occurred more recently (Figure 5d). This did not occur in the network without Hebbian learning, where performance for target 1 was worse than that for other targets despite the larger amount of practice (*p* = 0.011). This result shows that, in this trained network model as in the single-neuron case, associative Hebbian learning enables practiced behaviors to be better retained, protecting them from being overwritten as new behaviors are learned.

## Discussion

In this work, we have developed a bottom-up, mechanistic theory of fast and slow learning in parallel descending pathways, applying this theory to obtain insight into the possible roles of cortical and thalamic inputs to the basal ganglia in the acquisition and retention of learned behaviors through practice. While two-pathway architectures have been proposed previously as a means of addressing multistage learning in mammalian [48, 49] and songbird [50] motor circuits, those models featured different learning mechanisms and were applied to different anatomical structures than the two-pathway model studied here. In particular, our theory proposes two simultaneously occurring learning processes: a fast process that minimizes errors in the cortical pathway, paired with a slow process that strengthens associations in the thalamic pathway. From a computational perspective, this work provides a quantitative theoretical framework in which the effects of learning, forgetting, and practice can be analyzed for the simplest possible case, namely a single neuron classifying random input patterns. From a cognitive perspective, the work provides an explanation for why repeated practice can continue to have effects even after a memory or behavior has been learned to be performed with zero error. From a neurobiological perspective, this work provides a framework for understanding the automatization of learned behavior and the formation of habits, proposing how these processes might be implemented in neural circuits.

In the Introduction, we described the multiple effects of practice: improved performance, habit formation, and decrease in cognitive effort [3]. The two-pathway model—and its proposed implementation in the brain’s sensorimotor circuitry in particular—provides a unified account of these effects. First, fast learning in the corticostriatal pathway provides a mechanism for improved performance, with the greatest gains coming in the early stages of practice. Second, the two-pathway model provides a mechanistic description of habit formation, in which behaviors are produced consistently in response to particular stimuli, even when the behavior is no longer rewarded. This is because information about rewards and errors is encoded in the updates to corticostriatal synapses, which play less and less of a role in driving downstream activity the more that a behavior is practiced. Going further, the model may even provide a description of addiction and compulsion, in which behaviors persist even if they lead to negative outcomes. Finally, the two-pathway model provides a mechanistic description of behavioral automatization, with the amount of conscious effort devoted to a behavior decreasing with increasing amounts of practice, as control of the behavior is transferred from a cortical to a subcortical circuit.

While the model that we have developed was constructed to address procedural learning of motor behaviors, the results shown in Figure 2c-e suggest that it may also lead to insights for declarative memory. Not all of the results obtained from the two-pathway model were in agreement with published results from experiments on human memory, however. The fact that memory recall performance decays roughly as a power law has been relatively well established experimentally [45]. However, the forgetting curves in the two-pathway model are better described by exponential decay than by power laws (Supplemental Figure 4). In addition, experimental study of spaced-repetition effects in human memory has shown that there exists an optimal interval between presentations during training [51], such that the interval between the two training presentations should match that between the second presentation and the time of testing. In the two-pathway model, however, we found that shorter training intervals always lead to better testing performance, regardless of the testing interval (Supplemental Figure 5). Of course, it is not surprising that a simple single-neuron model fails to capture the full range of experimental results on human memory. Characterizing the additional mechanisms that could lead to such effects as power law memory decay and optimal intervals for spaced repetition will be a promising direction for future study.

A promising direction for future work would be to extend to two-pathway model in ways that might enable it to address effects such as power-law memory decay and spaced repetition. For example, one successful approach for obtaining power law memory decay has been to allow individual synapses to have multiple states, with a different timescale associated with each state [52, 39], and incorporating associative learning and the effects of practice into such multi-state models could be a promising direction for future work on the mechanistic modeling of learning and memory. More broadly, future work in this area will need to identify the network-level mechanisms that emerge in circuits with large numbers of neurons and multiple layers and might underlie phenomena such as power-law forgetting curves and the effects of spaced repetition. In addition, memory performance can also be affected by higher-level cognitive strategies such as chunking or creating narratives to link information meaningfully [45]. Clearly, such complex strategies cannot be addressed by a single-neuron model, though it might be hoped that such a model could be a building block for addressing such phenomena in the future.

Future work should also extend the model from supervised learning to reinforcement learning, which is less theoretically tractable but more plausible as a model of reward-based learning at corticostriatal synapses [14]. Reinforcement learning is also more plausible from a behavioral perspective, since it allows for performance to improve gradually by trial and error, leading to a richer description of learned motor behaviors than the single-step perceptron learning rule can provide. While a first step in this direction was taken in Supplemental Figure 6, which shows that the mechanisms of input alignment and control transfer are realized in a circuit trained with reinforcement learning, extending the theoretical framework underlying the two-pathway model to the case of reinforcement learning is an important direction for future work. Finally, in this work we have focused on sensorimotor striatum and its inputs. The thalamocortico-basal ganglia circuits that are involved in higher-level cognitive functions and that exist in parallel with the sensorimotor circuit could provide alternative interpretations of the model that we have presented [12, 53]. More broadly, this two-pathway motif might be found in other brain areas and could in part explain the large degree of apparent redundancy found throughout the neural circuitry of the brain.

## Acknowledgments

The authors are grateful to B.P. Ölveczky, members of the Ölveczky lab, and L.F. Abbott for helpful discussions and feedback. Support for this work was provided by the National Science Foundation NeuroNex program (DBI-1707398), the National Institutes of Health (R01 NS105349, DP5 OD019897, and U19 NS104649), the Leon Levy Foundation, and the Gatsby Charitable Foundation.

## Materials and Methods

Parameters and equations for the simulations in Figures 1, 2, 3, and 4, as well as Supplemental Figures 2 and 3, are provided in the Main Text.

For the simulations in Figure 5, *n* = 21 networks were trained for each of the conditions shown (SL only, SL+HL, and SL shuffled), where each network had *N*_*x*_ = *N*_*y*_ = 50, *N*_*z*_ = 10, and a 2-dimensional linear readout. The readout controlled the velocity of a cursor, which began at position **r** = **0** and moved to one of the four target positions: (1,0), (0,1), (−1,0), (0,-1). The target velocity in each trial was given by

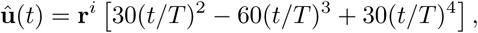

where *t* is the timestep, *T* = 10 is the number of timesteps in each trial, and **r**^*i*^ denotes one of the four target positions. This bell-shaped velocity profile describes a minimum-jerk trajectory from the initial to the target point. Both input layer populations receive fixed random projections of the network readout from the previous timestep, as well as a tonic random input associated with one of the four targets.

During training, the mean squared difference between the network output and **û**(*t*) was minimized by gradient descent on the weights **W** (labelled SL in Figure 5b), with learning rate *η*_SL_ = 0.001 (found by grid search). The weights **V** (labelled HL in Figure 5b) were either not trained (in the cases SL only and SL shuffled) or trained with the Hebbian update rule Δ*V*_*ij*_ = *η*_HL_[−*V*_*ij*_ + *z*_*i*_(*t*)*y*_*j*_(*t*)], where *η*_HL_ = 10^−6^ was found by grid search. The networks were sequentially trained on each target, as shown in Figure 5c. The bars in Figure 5d show the median test loss over the trained networks for each target, with error bars denoting the 25th and 75th percentiles.

## Appendix 1: The perceptron forgetting curve

As described in the main text, we wish to train a perceptron having *N*_*x*_ = *N* inputs and subject to the update rule (1) to map *P* random input patterns onto randomly chosen binary outputs. The question that we seek to answer is, after *P* patterns have been trained using the update rule in (1), what will be the probability of misclassification if we then test the output produced by a particular pattern *ν* without any further learning? Clearly, the most recently learned patterns are likely to produce correct outputs, while those learned long in the past are more likely to produce errors due to accumulated changes in the weights **w** during subsequent learning. In general, the probability of an error when testing on pattern *ν* is, using the Heaviside step function Θ(·), given by

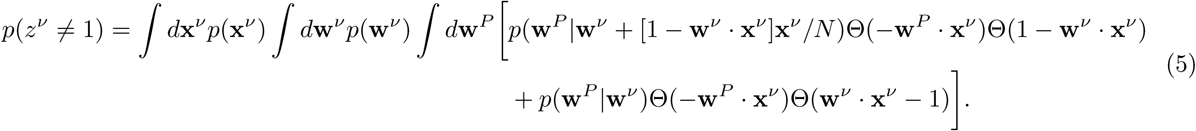

In this equation, we have, without loss of generality, redefined 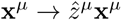, so that the target output becomes *z*^*μ*^ = 1 for every pattern. We have also set the classification margin *κ* = 1, which amounts to a choice for scaling the overall magnitude |**w**|. In this equation, the weight vector just before training pattern *ν* is assumed to come from a distribution *p*(**w**^*ν*^), which we shall derive below. The first line in (5) counts the cases in which the classification using this weight vector is initially incorrect or correct with margin less than *κ* = 1 (so 1 − **w**^*ν*^ · **x** *>* 0) and in which, after making the initial update **w**^*ν*^ → **w**^*ν*^ + Δ**w**^*ν*^, the weight vector evolves through *P* − *ν* successive updates into the final weight vector **w**^*P*^, which leads to incorrect classification when pattern *ν* is again tested (−**w**^*P*^ · **x**^*ν*^ *>* 0). Similarly, the second line of (5) counts the cases in which the classification is initially *correct* with a sufficiently large margin and in which, after making successive weight updates, the final weight vector **w**^*P*^ again leads to incorrect classification.

At this stage, the probability distributions *p*(**w**^*ν*^) and *p*(**w**^*P*^ |**w**^*ν*^) in (5) are unknown. For the latter distribution, however, we can track its evolution step by step using the update rule (1). Because **x** is a random variable, we first seek to find the distribution *p*(Δ**w**|**w**) by averaging over **x**. In fact, it will be sufficient just to calculate the first two moments of this distribution. Let **x** = **x**^‖^ + **x**^⊥^, where **x**^‖^ is the component along **w**, and the index *ν* has been dropped for simplicity. In this case the weight update (1), in the case where an update occurs, can be written as

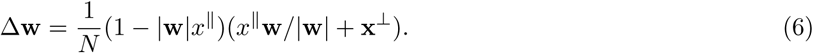

Using (6), in the case where a weight update occurs, the first moment is given by

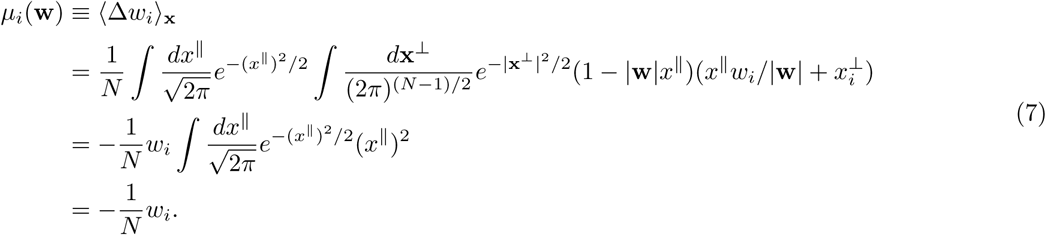

Similarly, the second moment of the distribution is

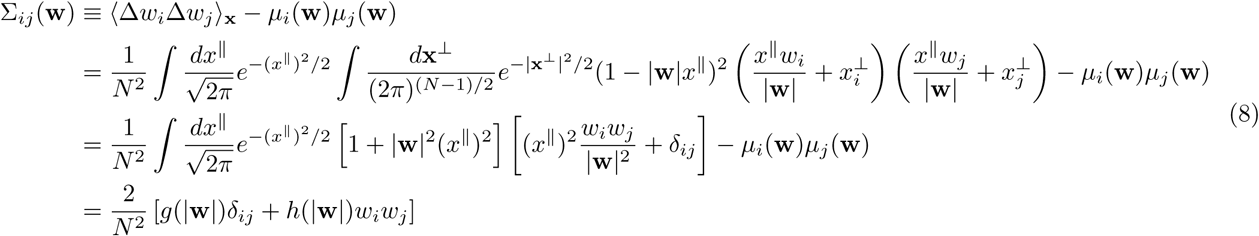

where

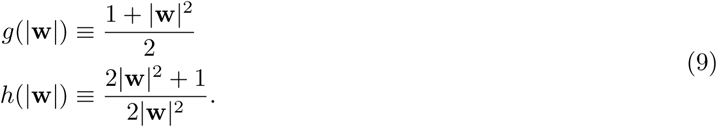

Together, (7)-(9) describe a drift-diffusion process. If we neglect higher-order moments, then the single-step probability distribution is given by

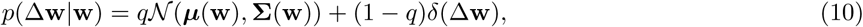

where *q* (to be calculated below) is the probability that a weight update occurs in a given step, and 𝒩 (·, ·) is the multinormal distribution. The time evolution of the probability distribution of the weights is then given by the Fokker-Planck equation [54]:

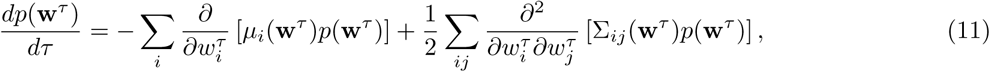

where the initial condition from (5) is either *p*(**w**) = *δ*(**w**−**w**^*ν*^) or *p*(**w**) = *δ*(**w**−**w**^*ν*^ −Δ**w**^*ν*^), and *τ* ≡ *q*(*P* −*ν*) is the effective time variable.

Because the coefficients in (11) depend on |**w**^*τ*^|, the full solution is not known in general. However, it is straightforward to calculate the time evolution of the moments of *p*(**w**^*τ*^) by multiplying both sides by powers of **w**^*τ*^ and using integration by parts [54]. Denoting the first two moments (not to be confused with the moments of Δ**w** defined above) as

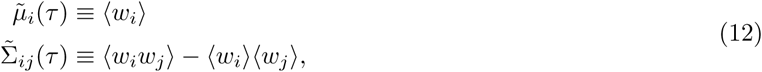

the time evolution is given by

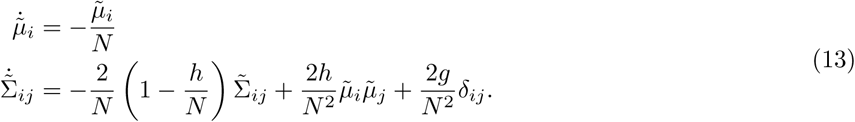

In the *N* → ∞ limit, we can assume (to be checked below) that |**w**^*τ*^ | is constant. In this case, (13) has the solutions

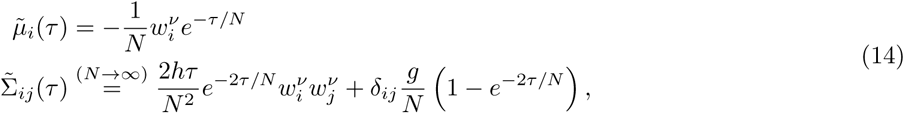

where terms ∼ *O*(1/*N* ^2^) in the second equation have been dropped in the large-*N* limit (while the first term is kept because *τ* may be ∼ *O*(*N*)).

With (14), the solution to the Fokker-Planck equation (11) when just the first two moments are kept is

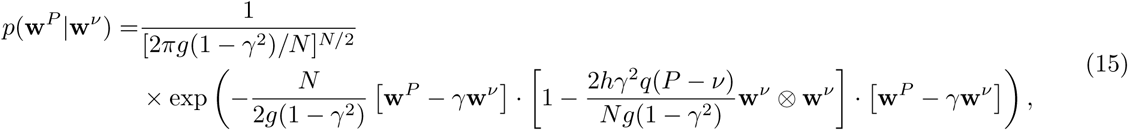

where “⊗” denotes the outer product (i.e. [**w** ⊗ **w**]_*ij*_ = *w*_*i*_*w*_*j*_), and we have defined

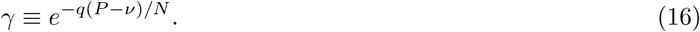

In the *N* → ∞ limit, the anisotropic term can be ignored, giving

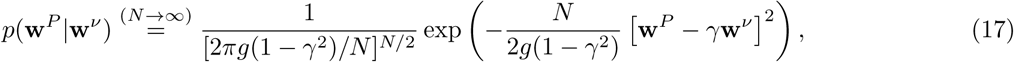

which is the probability density evolution corresponding to an Ornstein-Uhlenbeck stochastic process [54]. After a long time, *γ* → 0 and *p*(**w**^*P*^ |**w**^*ν*^) from (15) approaches the steady-state distribution

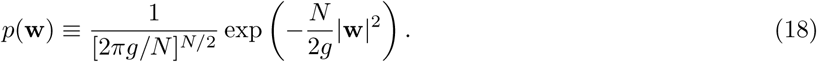

Thus, the deterministic update rule (1) leads to a bounded steady-state weight distribution *p*(**w**) after a large number of classifications have been learned. This differs somewhat from previous models of sequential learning with random synaptic weight updates, such a bounded distribution as *P*/*N* → ∞ was achieved either by requiring that the synaptic weights should be bounded [38] or that that they should decay slightly at each step [39].

Given the steady-state distribution (18), we can assume that *ŵ* ≡ |**w**| is constant in the large-*N* limit and let *ĝ* ≡ *g*(*ŵ*). Then, with *g* = *ĝ*, (18) can be used to calculate the variance of **w**, leading to the self-consistent equation *ŵ*^2^ = *ĝ*, which, using the definition of *g*(|**w**|) from (9), has the solution *ŵ*^2^ = *ĝ* = 1. This is close to but differs somewhat from the steady-state norm found in numerical simulations, from which *ŵ* ∼ 1.19. The reason for this is presumably because of the decision to approximate *p*(Δ**w**|**w**) using only the first two moments of the distribution in (11). In general, the higher-order moments do not vanish, and these will contribute higher-order derivative terms in the Fokker-Planck equation (12). In turn, such terms will lead to nonvanishing higher-order moments in the distribution *p*(**w**), beyond the two that were calculated in (12)-(14). Presumably, it is these higher-order terms which cause the discrepancy between the simulated result and the self-consistent calculation. In the theoretical curves shown in the Results section, we use the value of *ŵ* obtained from simulations. Using (18), we can also calculate the probability of making a weight update in a given step, which is given by

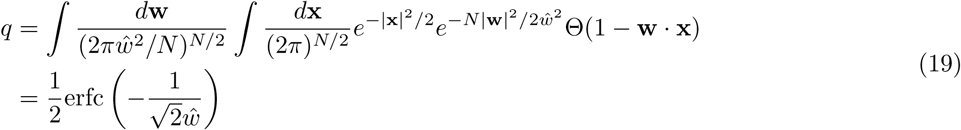

and evaluates to *q* ≃ 0.798.

With the preceding points in mind, and making use of the probability distributions (17) and (18), we can proceed to evaluate the integrals in (5) to obtain the probability of incorrect classification when testing pattern *ν*. In order to factorize the arguments of the Heaviside step functions, we make use of the following identity:

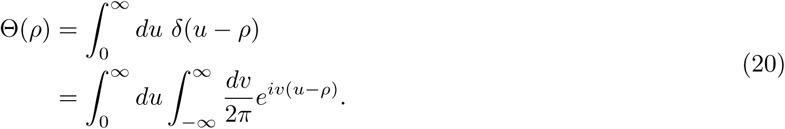

with this trick, (5) becomes

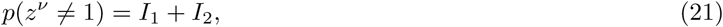

where

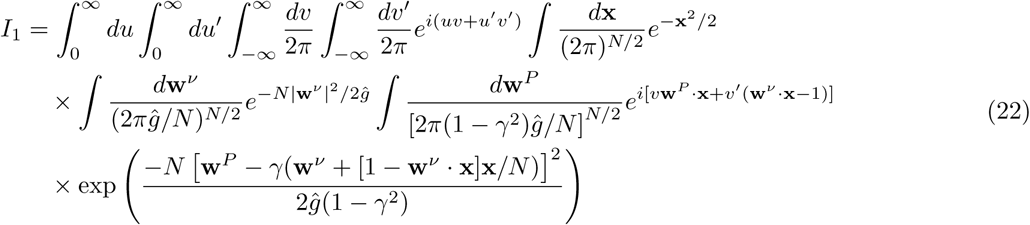

and

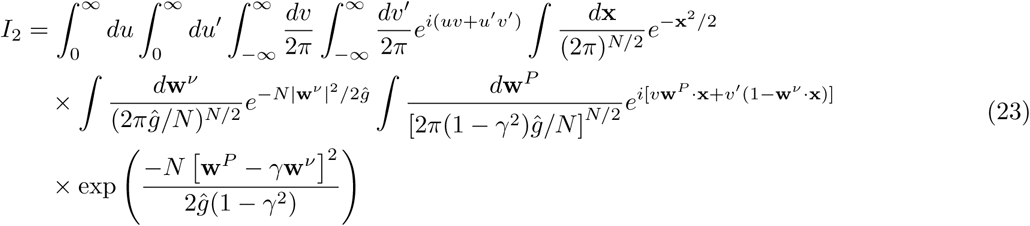

Beginning with *I*_1_, the integral over **w**^*P*^ can be performed, which, after simplification and using **x**^2^ = *N* in the *N* → ∞ limit, leads to

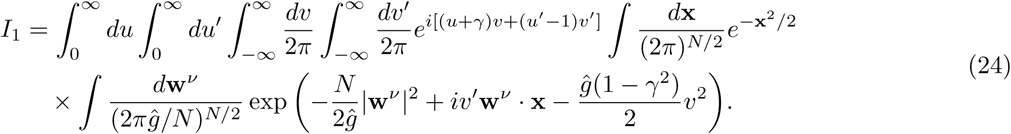

Performing the integrals over **w**^*ν*^ and **x** then leads to

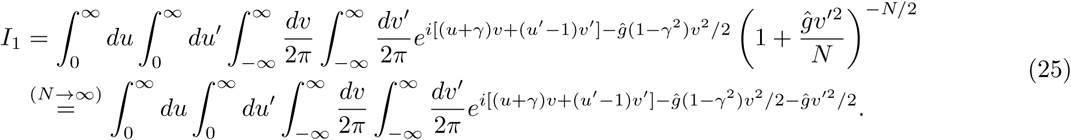

Finally, the integrals over *v* and *v*′ can be performed exactly, then those over *u* and *u*′ can be evaluated using the complementary error function, yielding the final result

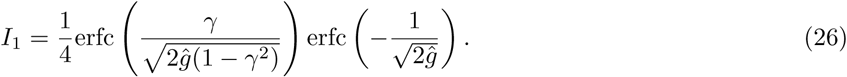

In a similar manner, the integrals in (23) can be evaluated to get

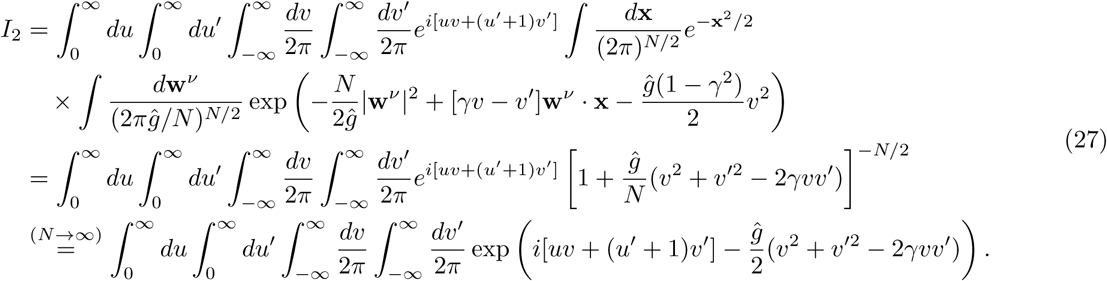

After performing the integrals over *v, v*′, and *u*′, then changing variables for the *u* integral, the final result is

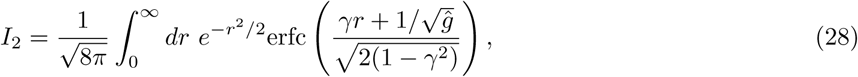

where the final integral in this case must be performed numerically.

As a check, we can evaluate these results in their extreme limits. In the case *γ* = 1, which corresponds to testing the most recently learned pattern, we have *I*_1_ ∼ *I*_2_ ∼ erfc(∞) → 0, so that there is perfect classification for very recently learned patterns. In the opposite limit of *γ* = 0, which corresponds to testing patterns learned in the distant past, we obtain 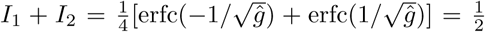, which means that very old patterns are completely overwritten and so are classified at chance level.

## Appendix 2: The two-pathway forgetting curve

Let us introduce a second source of input to the downstream units, so that *z*^*μ*^ = *φ*(**w** · **x**^*μ*^ + **v** · **y**^*μ*^), where 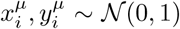. Though it is not necessary, we will assume for notational simplicity that the numbers of units in the two input layers are the same, so that *N*_*x*_ = *N*_*y*_ = *N*. The weights **w** are again trained using supervised learning:

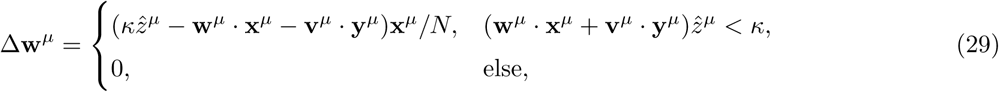

The second set of weights, meanwhile, is updated using the following rule:

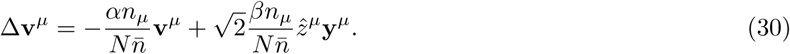

This Hebbian update rule defines an *N* -dimensional Ornstein-Uhlenbeck stochastic process with time-dependent coefficients. The evolution of the probability distribution *p*(**v**, Δ*t*) is given by the master equation:

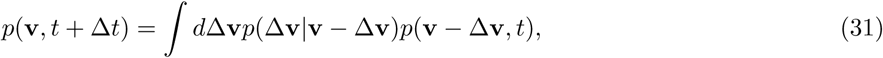

where

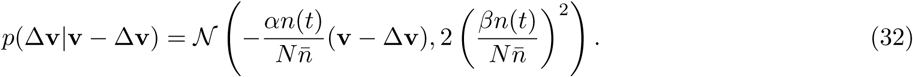

We can further note that, since the components of **v** are not coupled to one another in (30), we can, without loss of generality, consider the evolution of just a single component *v*_*i*_. In this case the two sides of (31) can be expanded to obtain

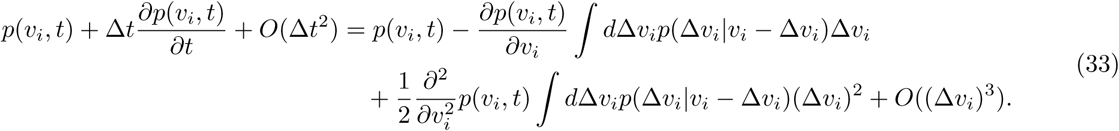

Then, using (32) to obtain ⟨Δ*v*_*i*_⟩ and ⟨ (Δ*v*_*i*_)^2^⟩, (33) leads to the Fokker-Planck equation, which describes the evolution of the probability distribution *p*(**v**) as new patterns are learned:

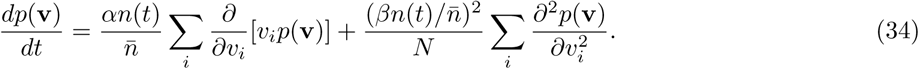

In this equation, we have taken the continuous-time limit by letting Δ*t* = 1/*N* and *t* ≡ (*μ* − *ν*)/*N*, where *μ > ν*, the number of repetitions to be *n*(*t*) ≡ *n*_*μ*_, and the initial condition to be given by the distribution (to be calculated below) *p*(**v**^*ν*^).

The Fokker-Planck equation (34) can be solved using the Fourier transform [54]

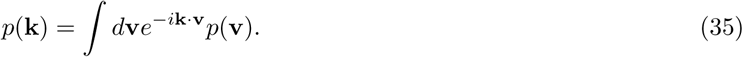

With this, (34) becomes

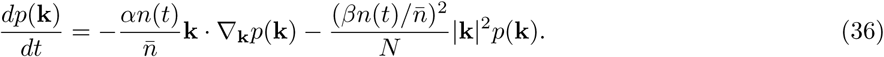

This equation can be solved by making the following ansatz:

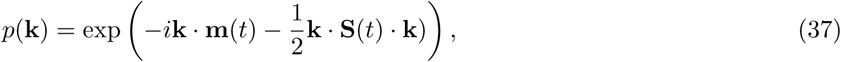

which, by substituting into (36) and requiring that the terms at each order in **k** vanish, leads to the following equations for the time-dependent coefficients:

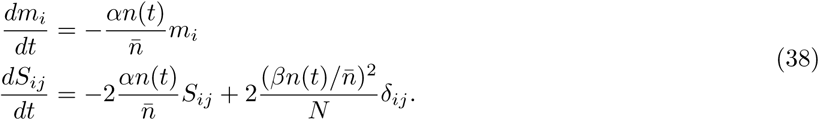

These equations then have the solutions

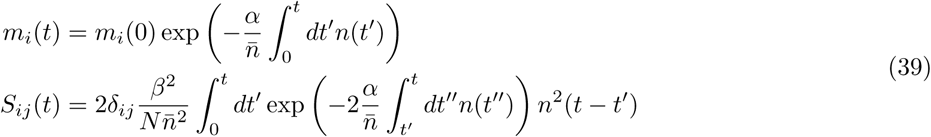

With this, and letting *t* = (*P* − *μ*)/*N*, we can take the inverse Fourier transform of (37) to obtain the distribution of the final weight vector given the weights at pattern *ν*:

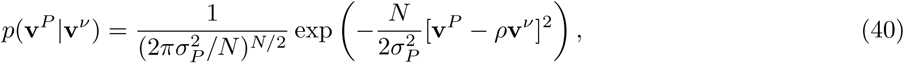

where we have identified 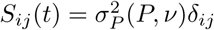, with

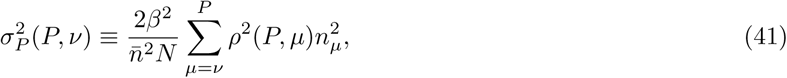

and we have also identified 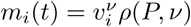, with

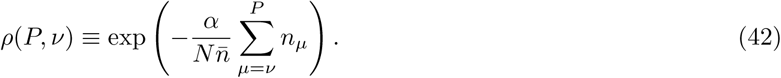

In what follows below, in order to keep expressions compact, we shall write 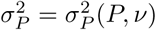 and *ρ* = *ρ*(*P, ν*). Further, though it is not strictly necessary, these expressions can be considerably simplified if we make the simplifying assumption that the average of *n*_*μ*_ in (42) over the last *P* − *ν* patterns is equal to its average 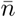 over the full set of *P* patterns (or, in the case of (41), that 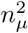 can be replaced by 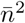). In this case, (42) becomes *ρ*(*P, ν*) = *e*^−*α*(*P*−*ν*)/*N*^, while (41), after performing the summation, becomes 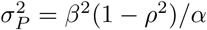.

Returning to the distribution (40), we can see that it begins as a *δ* function at **v**^*ν*^ and, as more patterns are introduced and *ρ* → 0, evolves to the following steady-state distribution:

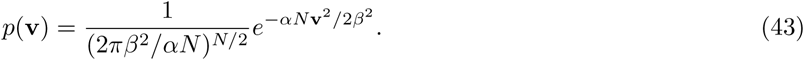

This is the distribution from which **v**^*ν*^ will be drawn in order to calculate the error rate for pattern *ν* below. In addition to driving a drift-diffusion process for the weights **v**, the updates to **v** also affect the evolution of the distribution of weights **w** due to the appearance of **v** in the update rule (29). In order to account for this change, the first two moments of the update Δ**w** must be reevaluated as in (7)-(8), but now averaging over the random variable **y** in addition to averaging over **x**. Because ⟨*y*_*i*_⟩ = 0, the first moment in (7) remains unchanged. As for the second moment, (8) generalizes to

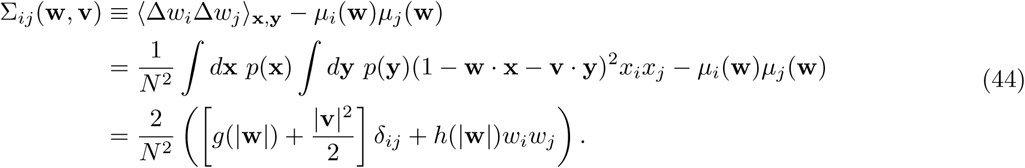

As before, we will assume that |**w**|^2^ and |**v**|^2^ can be replaced by their average values in the *N* → ∞ limit. Noting from (43) that ⟨**v**^2^⟩ = *β*^2^/*α*, we have the diffusion tensor 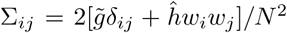, where 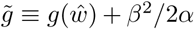, where, as before, 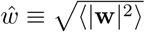 is taken from numerical simulations. From this result, we see that the equations (17)-(18) determining the evolution of **w** can also be applied in the two-pathway case by making the substitution 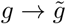.

As in the previous section, our goal is to calculate the probability of incorrect classification when testing pattern *ν* after training *P* patterns in sequence. Including the second input pathway, this quantity is given by

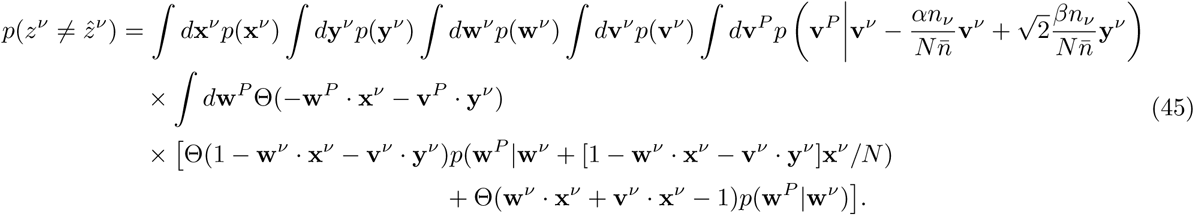

As before, we have absorbed all 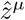 into the definition of the input activity vectors **x**^*μ*^ and **y**^*μ*^ by letting 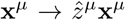 and 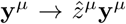, effectively setting all 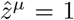. The first term in the integrand corresponds to cases in which the classification of pattern *ν* is initially incorrect, so that the weights **w** and **v** are both updated. The second term corresponds to cases in which the initial classification is correct, so that only the weights **v** are updated. In both cases, the weight distributions for **w** and **v** evolve according to drift-diffusion processes, as new patterns are learned up until pattern *P*, at which time, the classification of pattern *ν* is tested using the final weights **w**^*P*^ and **v**^*P*^ (the first Θ function appearing in the integrand).

Again using the trick (20) to represent the Heaviside step functions, the two terms in (45) can be written as

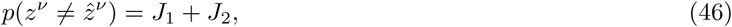

where

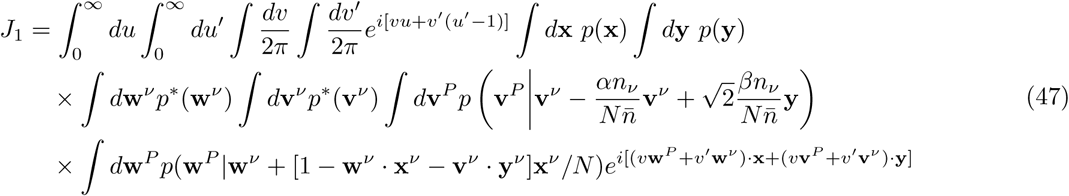

and

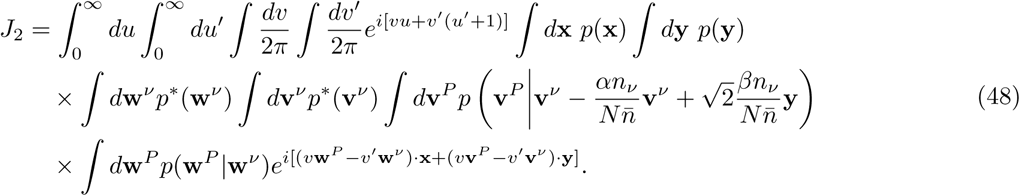

Beginning with *J*_1_, we can shift the integration variable **w**^*P*^ → **w**^*P*^ − *γ*(**v**^*ν*^ · **y**)**x***/N* and use **x**^2^ = *N* in the *N* → ∞ limit to obtain

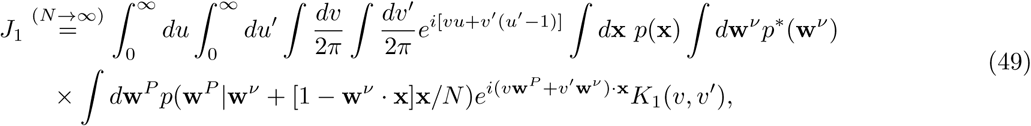

where

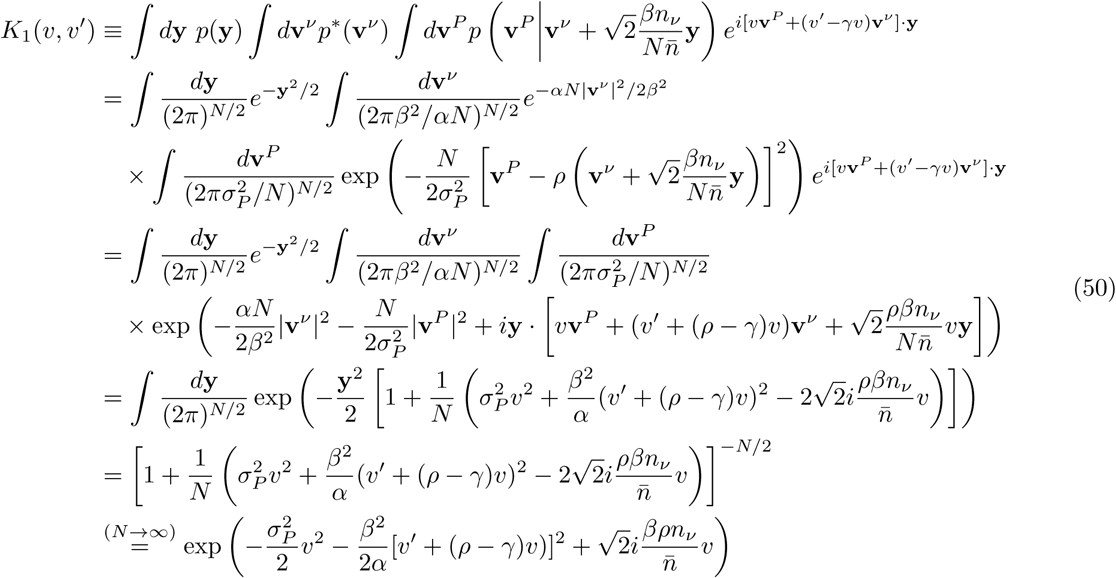

Where the Hebbian decay term ∼ *α***v**^*ν*^ */N* was dropped in the first line because it vanishes as *N* → ∞, and the integration variable **v**^*P*^ was shifted in the third line. Using this result, and noting that the integrals over **x, w**^*ν*^, and **w**^*P*^ are the same as those appearing in (22) (with 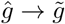), (49) becomes

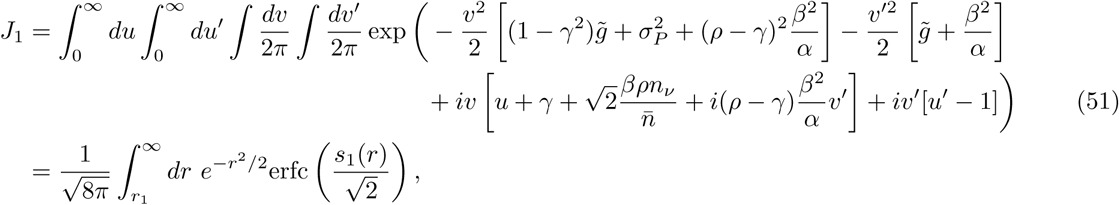

where we have defined

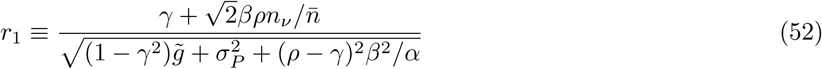

and

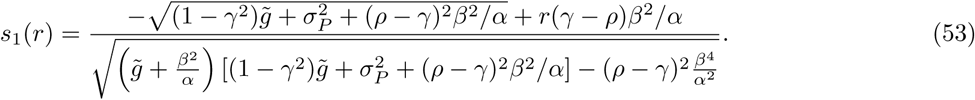

As in the case without the Hebbian pathway, the last remaining integral in (51) must be performed numerically.

Equation (48) for *J*_2_ can be evaluated in a similar manner. To begin, we express it as

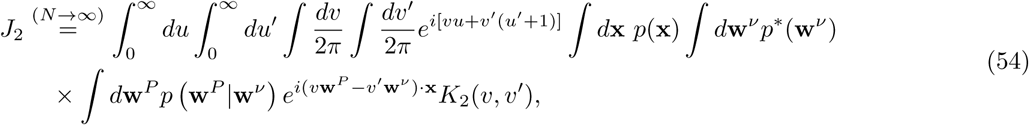

where

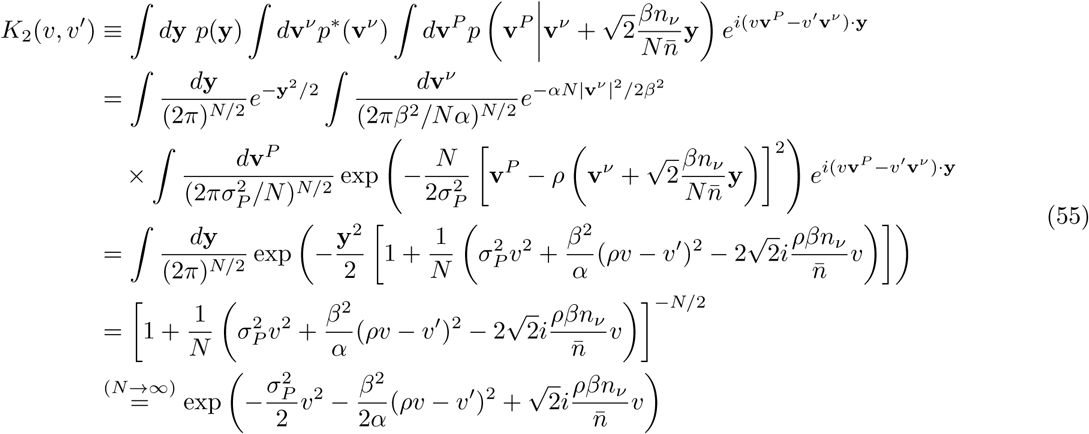

Using this result, and noting that the integrals over **x, w**^*ν*^, and **w**^*P*^ are the same as those appearing in (23) (with 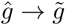), (54) becomes

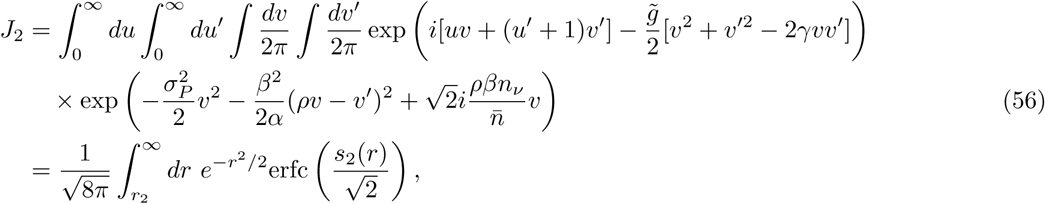

where we have defined

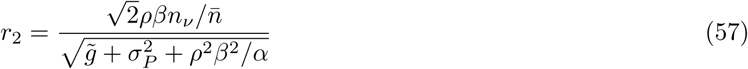

and

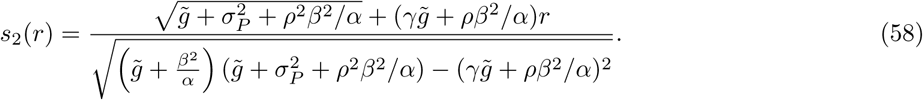

It is straightforward to check that, in the limit *β* → 0, which corresponds to shutting off the Hebbian input pathway, we recover *J*_1,2_ → *I*_1,2_ from the simple perceptron result.

Finally, we can calculate the probability of making a weight update in a given step, which in the two-pathway case is given by

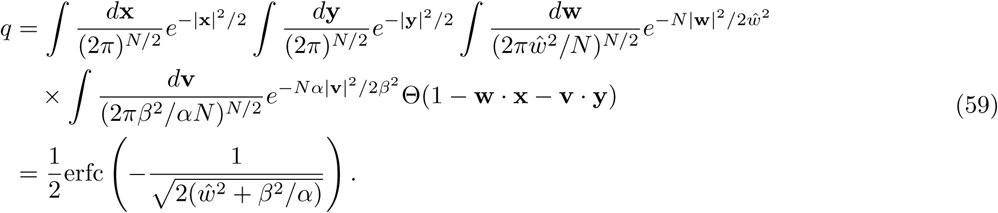

In the case where *β*^2^*/α* = 0, this evaluates to the earlier result *q* ≃ 0.798. For nonzero *β*, on the other hand, *q* decreases and approaches 0.5 as *β*^2^*/α* → ∞.

## Supplemental Information

### Learning with gradient descent

**Supplemental Figure 1:**
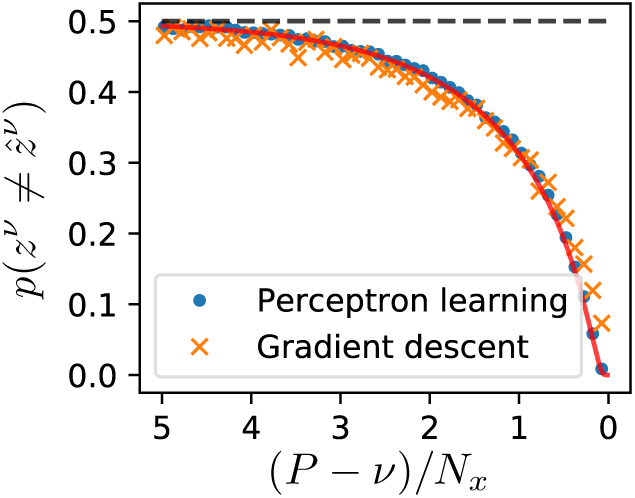
Training with gradient descent yields a forgetting curve similar to the perceptron case.

The perceptron learning rule, according to which synaptic weights are adjusted to produce the correct output in a single step, is mathematically convenient but biologically questionable. In order to address this, we simulated the forgetting curve using gradient descent learning, a widely used supervised learning algorithm in which small updates are accumulated over many repetitions to minimize the readout error.

In this case the output of the neuron is 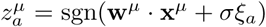, where *ξ*_*a*_ ∼ 𝒩(0, 1) is drawn randomly for step *a*. The number of steps for each pattern was chosen to be *N*_steps_ = 100, and the learning rate *η* = 0.01 and the noise amplitude *σ* = 0.2 were chosen by grid search to maximize the area between the forgetting curve and chance performance. The noise is not strictly necessary in gradient descent learning, but is included here so that a finite classification margin will be obtained, as in the case of the perceptron learning rule (Figure 1b). In this case, the synaptic weights were updated according to the gradient-descent update rule 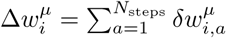, where

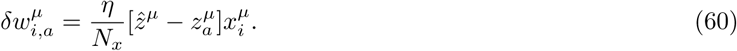

As shown in Supplemental Figure 1, the forgetting curve obtained for the perceptron with this alternative learning rule is similar to that obtained using the perceptron learning rule.

### Two-pathway forgetting curves depend on Hebbian learning and decay rates

**Supplemental Figure 2:**
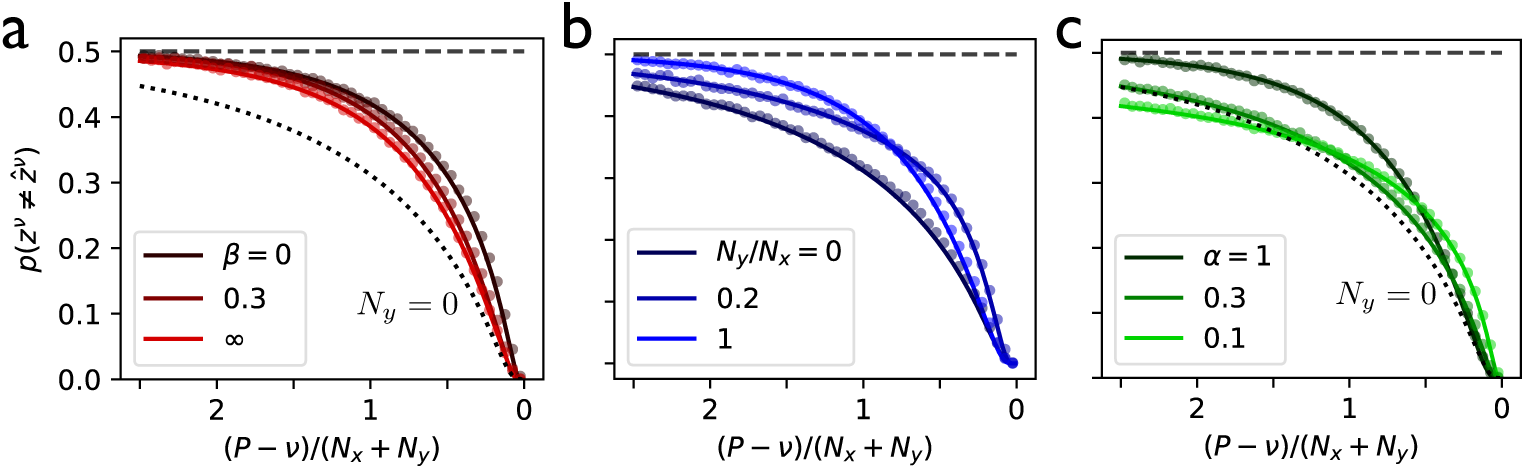
Forgetting curves in the two-pathway model with each pattern trained once. **(a)** The forgetting curve for different values of the Hebbian learning rate *β*, with *α* = 1 and *N*_*x*_ = *N*_*y*_ (dotted line shows the case with no second pathway). **(b)** The forgetting curve for different values of *N*_*y*_ */N*_*x*_, with *α* = *β* = 1. **(c)** The forgetting curve for different values of the Hebbian weight decay rate *α*, with *β* = 1 and *N*_*x*_ = *N*_*y*_. In (b)-(d), solid lines are theoretical results; points are simulations with *N*_*x*_ = *N*_*y*_ = 1000.

In the two-pathway model, we kept 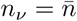 constant for all patterns and investigated the dependence of the forgetting curve (4) on its other parameters. Starting with the Hebbian learning rate, we found that nonzero values of *β* shifted the forgetting curve slightly downward, modestly reducing the error rate for all patterns (Figure 2b). Whether this qualifies as a true improvement, however, depends somewhat on bookkeeping. For a fixed total number of synapses *N*_*x*_ + *N*_*y*_, the error rate can be reduced by allowing for Hebbian learning. However, the error rate is reduced even more by eliminating the Hebbian synapses entirely (dashed curve in Figure 2b), which decreases the denominator *N*_*x*_ + *N*_*y*_ by setting *N*_*y*_ = 0. Stated differently, if the goal is to minimize the error rate for a fixed total number of synapses, this is accomplished most effectively by letting all of the synapses be updated with supervised learning rather than with Hebbian learning (Figure 2c). If, on the other hand, the goal is to minimize the error rate for a fixed number *N*_*x*_ of supervised synapses, then a benefit is obtained from including additional synapses with Hebbian learning. The exception to these conclusions occurs for small values of the Hebbian decay rate *α*, in which case very old memories can persist for longer with error rates below chance level (Figure 2d). In this case, there is a benefit to adding Hebbian synapses even if they are counted against the total *N*_*x*_ + *N*_*y*_.

### Optimal forgetting rate and memory capacity

**Supplemental Figure 3:**
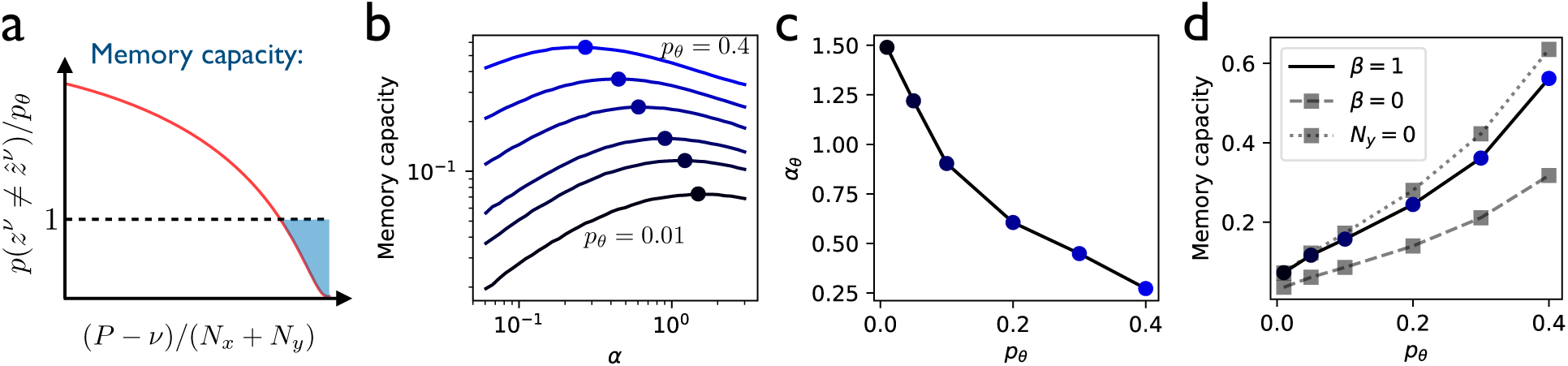
Error tolerance determines the optimal forgetting rate and memory capacity. **(a)** Given a threshold *p*_*θ*_ of incorrect classification probability, the memory capacity is defined as the area above the forgetting curve and below the threshold, normalized by *p*_*θ*_. **(b)** For each value of *p*_*θ*_, the memory capacity is optimized with respect to the Hebbian forgetting rate *α*. (For all curves, *β* = 1 is fixed, *N*_*x*_ = *N*_*y*_, and 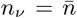 for all patterns.) **(c)** The optimal forgetting rate from (b) as a function of *p*_*θ*_. **(d)** The memory capacity from (b) as a function of the classification threshold *p*_*θ*_. Dashed line shows the capacity with a second pathway with no learning (*N*_*y*_ = *N*_*x*_, *β* = 0); Dotted line shows the capacity with no second pathway (*N*_*y*_ = 0).

We investigated the effects of the parameters *α* and *β* in the two-pathway model, while still holding *n*^*ν*^ constant for all patterns, by setting a threshold *p*_*θ*_ for the acceptable error rate, then defining the memory capacity as the integrated area between the forgetting curve and this threshold value, normalized by *p*_*θ*_ (Supplemental Figure 3a). Because the forgetting curve was found to have relatively weak dependence on *β* (Figure 2b), we fixed *β* = 1 and considered the effects of *α* and of *p*_*θ*_ on the memory capacity. For a given choice of *p*_*θ*_, we found the value of *α* that maximized the memory capacity (Supplemental Figure 3b). This led to the conclusion that, the larger the error tolerance *p*_*θ*_, the smaller the forgetting rate *α* should be in order to maximize the memory capacity (Supplemental Figure 3c). Finally, we found that the optimized memory capacity increases as the error tolerance becomes greater (Supplemental Figure 3d). Consistent with our observations from Figure 2, we found that the memory capacity, which is normalized by the total number of synapses *N*_*x*_ + *N*_*y*_, is improved by learning in the second pathway if *N*_*x*_ + *N*_*y*_ is held constant (solid vs. dashed curve in Supplemental Figure 3d), but is even larger if the second pathway is left out entirely (dotted curve in Supplemental Figure 3d), again indicating that adding supervised synapses is a better strategy than adding unsupervised synapses if the goal is to optimize the forgetting curve for a fixed total number of synapses.

### Forgetting curves decay exponentially

**Supplemental Figure 4:**
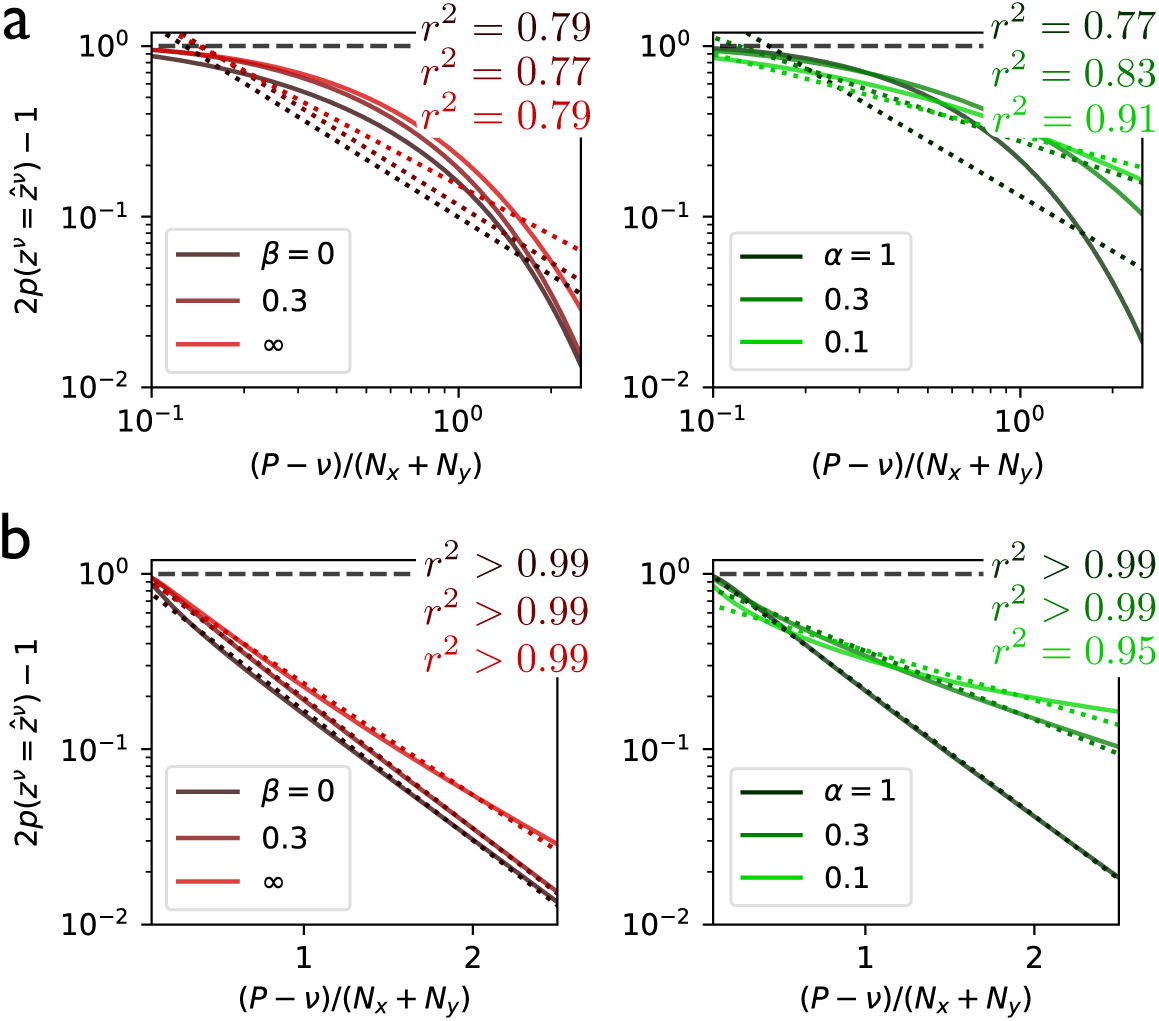
Forgetting curves exhibit approximately exponential decay. **(a)** Forgetting curves from Figure 2b (*left*) and 2d (*right*) plotted on logarithmic axes, with *N*_*x*_ = *N*. Dotted lines are best fits with power law decay. **(b)** Forgetting curves as in (a), but plotted with just one logarithmic axis. Dotted lines are best fits with exponential decay.

Many previous studies on memory have shown that forgetting curves are well described by curves decaying as a power law [45], in which the probability of correct recall has the form ∼ *t*^−*a*^, where *t* is time and *a* > 0. Supplemental Figure 4a shows that such a fitting function, which appears as a straight line in a log-log plot, does a relatively poor job of fitting the forgetting curves from the two-pathway model. In contrast, exponentially decaying functions, in which the probability of correct recall ∼ *e*^−*at*^ and which appear as straight lines on semilogarithmic plots, provide a good fit to the forgetting curves (Supplemental Figure 4b).

### Spaced repetition

**Supplemental Figure 5:**
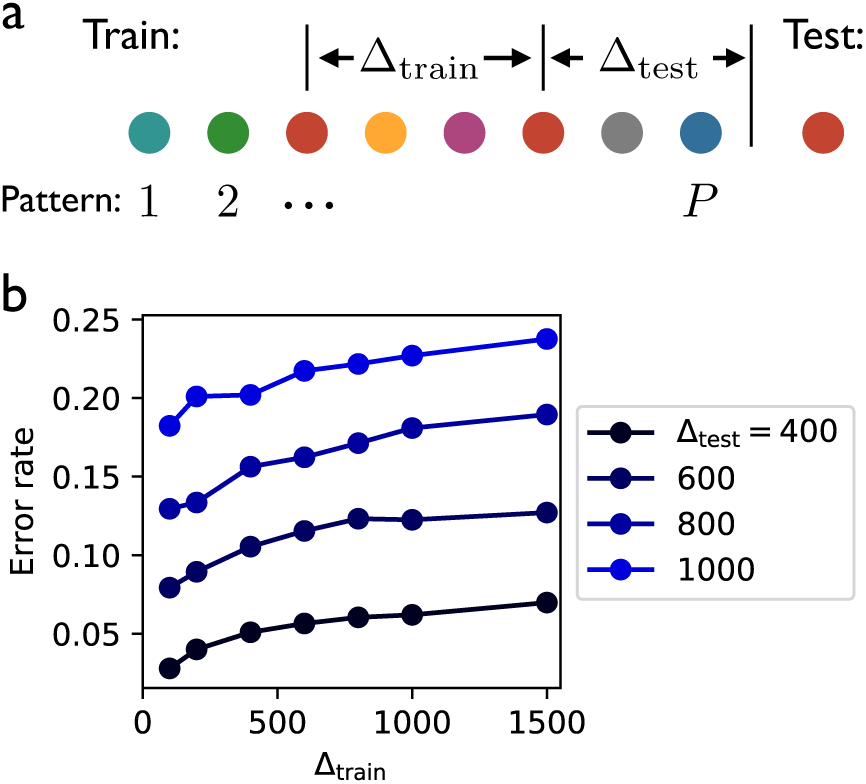
Short training intervals during spaced repetition lead to better testing performance. **(a)** The two-pathway network is trained with random sequential patterns, with one particular pattern presented twice. **(b)** For any interval Δ_test_ between the second presentation and the testing phase, the testing performance for the repeated pattern is best when the interval between the presentations during training is short. Results are simulations from a two-pathway network with *N*_*x*_ = *N*_*y*_ = 1000 and *α* = *β* = 1.

In simulations of the two-pathway model, the neuron was trained in sequence to perform *P* classifications. All of the input patterns were distinct, except for a single pattern that was presented twice during training, with an interval Δ_train_ between presentations. The interval between the second presentation and the testing phase was Δ_test_. For any Δ_test_, smaller values of Δ_train_ always led to a lower error rate for the repeated pattern during testing (Figure 5b).

In addition, we note that repetition of patterns has no significant effect at all in the single-pathway model without Hebbian weights. In this case, upon the second presentation of the repeated pattern during training, the weight vector will either not be updated at all (if classification is already correct with sufficient margin) or will be updated to lie on the correct side of the classification boundary plus a margin (if the classification is initially incorrect or correct with insufficient margin). In neither of these two cases is there a benefit to having seen the pattern before, since one of these same things would have happened if the first presentation had not occurred.

Thus, while the information accumulated in the Hebbian weights of the two-pathway model is an important ingredient for describing nontrivial effects due to spaced repetition, the single-neuron model appears to be unable to account for the experimentally observed existence of optimal repetition intervals [51], and thus leaves room for future work on this topic.

### Input alignment with reinforcement learning

**Supplemental Figure 6:**
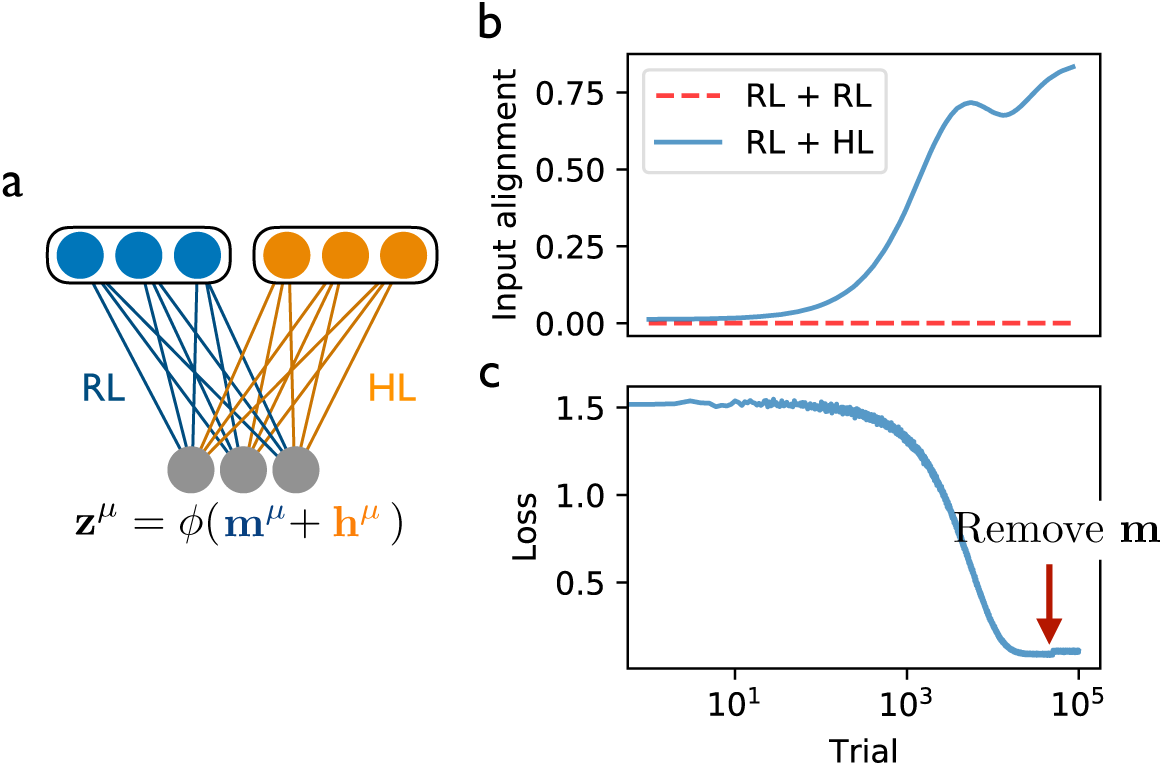
Input alignment and control transfer in a network with reinforcement learning and Hebbian learning. **(a)** Version of the two-pathway model in which synaptic weights in the first pathway are updated with RL. **(b)** As training is repeated, the normalized alignment between **m** and **h** grows. The alignment does not occur if the second input pathway is updated with RL instead of Hebbian learning (dashed line) **(c)** The mean squared error between the output activity and the target state decreases with training and remains low if the first input pathway is removed.

We checked that the mechanisms of input alignment and control transfer could also be obtained using reinforcement learning (RL), which is a more plausible form of synaptic plasticity than the perceptron learning rule for describing changes at corticostriatal synapses. In this case, a noise current *ξ* was added to the readout layer, with *ξ*_*i*_ ∼ 𝒩(0, 1), and the updates to input weights in the first pathway were given by [55, 56]

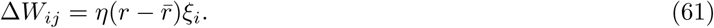

Over the course of many trials, where each trial consists of a complete presentation of all *P* = 10 patterns, the input currents from the two pathways become aligned when one pathway was trained with RL and the other with HL, but not if both pathways were trained with RL (Supplemental Figure 6b). Further, due to input alignment and control transfer, the first input **m** can be removed once the network is trained, without significantly impairing performance (Supplemental Figure 6c).

